# Characterization of Viral Insulins Reveals White Adipose Tissue Specific Effects in Mice

**DOI:** 10.1101/2020.08.21.261321

**Authors:** Martina Chrudinová, Francois Moreau, Hye Lim Noh, Terezie Páníková, Lenka Žáková, Randall H. Friedline, Francisco A. Valenzuela, Jason K. Kim, Jiří Jiráček, C. Ronald Kahn, Emrah Altindis

## Abstract

Members of the insulin/IGF superfamily are well conserved across the evolutionary tree. We recently showed that four viruses in the *Iridoviridae* family possess genes that encode proteins highly homologous to human insulin/IGF-1. Using chemically synthesized single chain (sc), i.e. IGF-1-like, forms of the viral insulin/IGF-1 like peptides (VILPs), we previously showed that they can stimulate human receptors. Because these peptides possess potential cleavage sites to form double chain (dc), i.e. more insulin-like, VILPs, in this study, we have characterized dc forms of VILPs for Grouper iridovirus (GIV), Singapore grouper iridovirus (SGIV) and Lymphocystis disease virus-1 (LCDV-1). GIV and SGIV dcVILPs bind to both isoforms of human insulin receptor (IR-A, IR-B) and to the IGF1R, and for the latter show higher affinity than human insulin. These dcVILPs stimulate IR and IGF1R phosphorylation and post-receptor signaling in vitro and in vivo. Both GIV and SGIV dcVILPs stimulate glucose uptake in mice. In vivo infusion experiments in awake mice revealed that while insulin (0.015 nmol/kg/min) and GIV dcVILP (0.75nmol/kg/min) stimulated a comparable glucose uptake in heart, skeletal muscle and brown adipose tissue, GIV dcVILP stimulated ~2 fold higher glucose uptake in white adipose tissue (WAT) compared to insulin. This was associated with increased Akt phosphorylation and glucose transporter type 4 (GLUT4) gene expression compared to insulin. Taken together, these results show that GIV and SGIV dcVILPs are active members of the insulin superfamily with unique characteristics. Elucidating the mechanism of tissue specificity for GIV dcVILP will help us to better understand insulin action, design new analogues that specifically target the tissues, and provide new insights into their potential role in disease.

## INTRODUCTION

In vertebrates, the insulin gene superfamily includes insulin, two insulin-like growth factors (IGF-1 and IGF-2), and more distant hormones including relaxin and the Leydig insulin-like peptides^1^. Insulin-like peptides have also been identified in invertebrates including insects, mollusks and nematodes^2–5^. These ligands are well conserved across the phylogenetic tree. However, their functions vary from the control of longevity and stress resistance in invertebrates to the control of metabolism and cell growth in vertebrates^5^. In mammals, insulin mainly regulates glucose and lipid metabolism^6^, whereas IGF-1 and IGF-2 control predominantly cell growth, proliferation and differentiation^7^. Insulin and IGF-1 bind to two different tyrosine kinase receptors in mammals – the insulin receptor, which itself exists in two isoforms (IR-A and IR-B), and the IGF-1 receptor (IGF1R)^7^. Invertebrates, on the other hand, often have multiple insulin-like peptides and elicit their biological function through one receptor or in rare cases through multiple receptors^2–4,8^. A major difference between insulin and IGF-1/2 is their ability to be processed post-transcriptional into either a two chain peptide hormone, in the case of insulin, or a single chain peptide hormone, in the case of IGF-1/2.

We recently showed that four viruses that belong to *Iridoviridae* family possess genes that show significant homology to human insulin/IGF-1 which we termed viral insulin/IGF-1-like peptides or VILPs for short^9,10^. Although viruses encoding these sequences were originally isolated from fish^11–14^, reanalyzing published human microbiome data, we identified the DNA of some of these viruses in human fecal and blood samples^9^. In our previous study, three VILPs were also chemically synthetized as single chain peptides (sc), i.e. IGF-1 like peptides, and we showed that scVILPs are weak ligands of the insulin receptor but strong ligands of the IGF-1 receptor in vitro and also possessed some glucose lowering effects in vivo^9^.

In humans, insulin is initially translated as a single chain peptide (preproinsulin) in pancreatic β-cells containing a signal peptide (SP) followed by B-, C- and A-domains (**Fig. 1**, **Fig. S1A**). Proinsulin is formed in endoplasmic reticulum by cleavage of the SP, and the C-peptide is removed in the secretory granules to form mature insulin with A- and B- chains bound together by disulfide bonds^15^. Unlike insulin, IGF-1 is produced in multiple tissues, but primarily in the liver^16^, and after cleavage of the signal peptide remains as a single chain peptide consisting of A- and B-domains, a short C-domain, and an additional D-domain at the C-terminus (**Fig. 1A**, **Fig. S1B**). The two peptide hormones show significant structural homology with about 50% of their amino acids being identical. Six cysteines that are crucial for the correct protein folding are also evolutionally conserved (**Fig. S1**). These cysteines are conserved throughout the phylogenetic tree^5,17,18^, and are considered to be a characteristic sign of the peptides belonging to the insulin/IGF peptide family. The main secondary structure motifs are also conserved between insulin and IGF-1, including the central α-helix in the B-chain/domain and two antiparallel α-helices in the A-chain/domain^19,20^.

**Figure 1:**
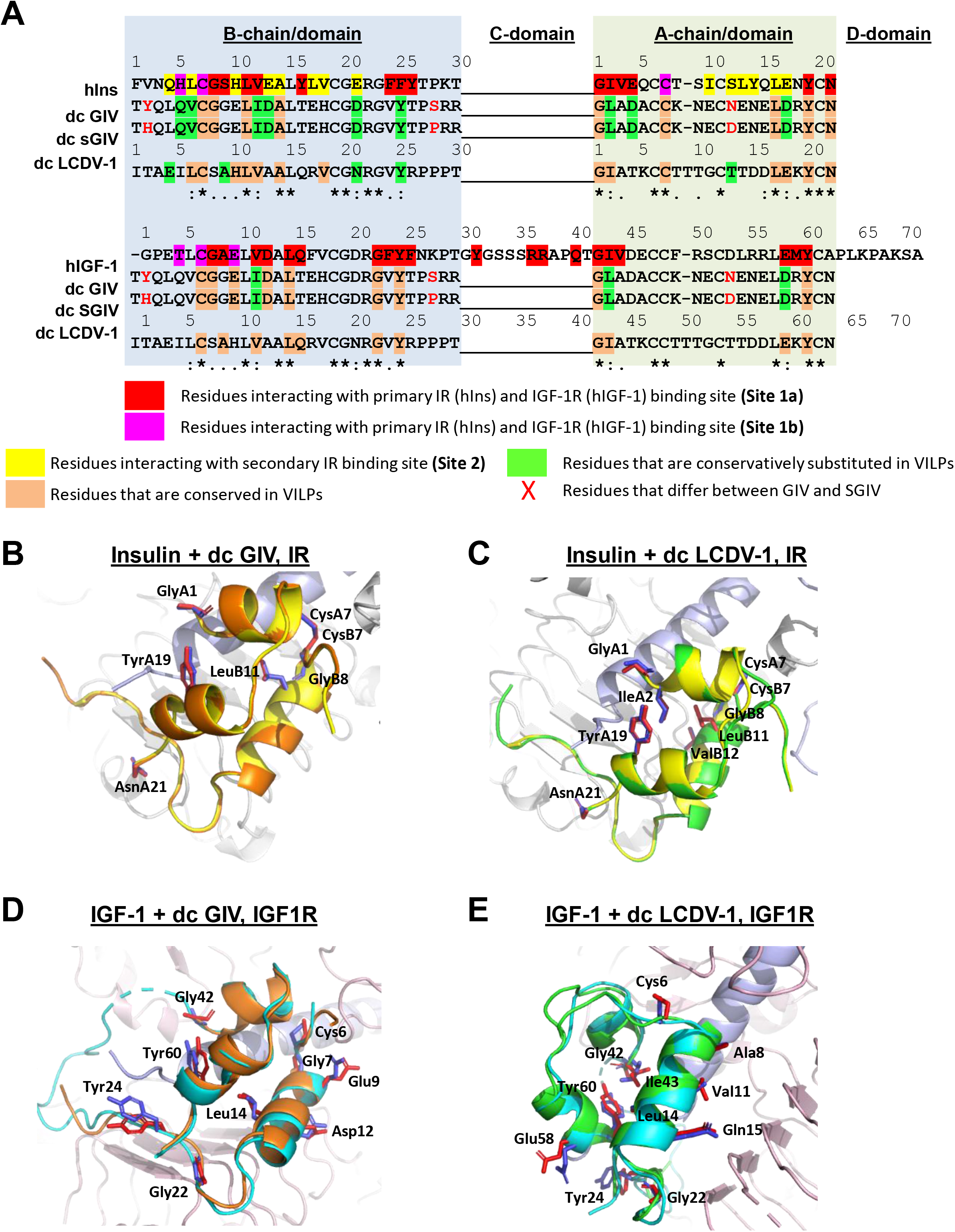
dcVILPs share significant homology in structure with human insulin and IGF-1. **A:** Sequence alignment of synthesized dcVILPs with human insulin and IGF-1. The residues important for receptor binding are highlighted with different colors. Site1a and Site1b are as described in^24^. **B - E:** Overlay of a model of GIV and LCDV-1 dcVILPs and insulin/IGF-1 bound to Site 1 of IR/IGF1R. Side chains of fully conserved amino acids are shown (main chain is shown for glycine). Insulin is in yellow, IGF-1 is in cyan, GIV dcVILP is in orange, LCDV-1 dcVILP is in green, IR is in grey, IGF1R is in pink.

In this study, we have synthesized and characterized the double chain (dc, insulin-like) forms of three VILPs - Grouper Iridovirus (GIV), Singapore Grouper Iridovirus (SGIV) and Lymphocystis disease virus-1 (LCDV-1) VILPs - for the first time. Using in vitro assays, we show that both GIV and SGIV dcVILPs can bind to both isoforms of the human insulin receptor and human IGF1R and stimulate post-receptor signaling, while LCDV-1 dcVILP is a very weak ligand. GIV and SGIV dcVILPs can also stimulate glucose uptake in vivo. During in vivo infusion experiments in awake mice, GIV dcVILP preferentially stimulates glucose uptake in white adipose tissue (WAT) compared to other insulin sensitive tissues. This is associated with higher Akt phosphorylation and expression of GLUT4 gene upon GIV dcVILP stimulation in WAT compared to insulin. Taken together, our results show that GIV and SGIV dcVILPs are potent members of insulin/IGF family and GIV dcVILP has unique WAT specific characteristics.

## RESULTS

### dcVILPs show significant homology in primary and predicted 3D structures with human insulin and IGF-1

Comparative alignment analysis revealed that GIV, SGIV and LCDV-1 dcVILPs show significant homology with human insulin and IGF-1 (**Table 1**. **Fig. S1**). All VILPs carry the six cysteine residues that form intrachain and interchain disulfide bonds and are critical for correct folding of insulin/IGF-like molecules (**Fig. S1**). While GIV and SGIV dcVILPs only differ in three amino acids within their A- and B-chains, the similarity between LCDV-1 and GIV/SGIV dcVILPs is lower with LCDV-1 sharing only 40% of amino acids of the A- and B-chains with GIV and SGIV dcVILPs (**Fig. S1**). A significant number of the residues that have been previously shown to be involved in insulin:IR or IGF-1:IGF1R interaction^21–26^ are either conserved or conservatively substituted in dcVILPs (**Fig. 1A**, **Table 1**). Structural studies have suggested that the mature IR or IGF1R can bind with high affinity two insulin or IGF-1 molecules effectively crosslinking two binding subsites on IR/IGF1R named as Site 1 and Site 2^21–23,27–29^. However, two recent studies suggested that up to four insulin molecules can bind to IR^24,25^ via two distinct binding sites. Uchikawa et. al.^24^ named the new binding site as “Site 2”, while they named previously identified sites as Site 1. In this manuscript, we used this new Site 1 and Site 2 nomenclature for IR binding. Interestingly, no analogic binding site to the new IR site 2 was identified in IGF1R^26^. Significant number of residues that are critical in receptor binding are conserved or conservatively substituted in dcVILPs (**Fig. 1A**, **Table 1**). To explore the similarity of 3D structures of dcVILPs with insulin, IGF-1 and its potential effect on binding to IR and IGF1R, we created models of dcVILPs bound to these receptors. These models indicate that residues conserved among dcVILPs, insulin and IGF-1 take similar positions upon receptor binding. Insulin and IGF-1 bound to Site 1 of IR/IGF1R, respectively, and the predicted structures of GIV and LCDV-1 dcVILPs bound to Site1 of the receptors are shown in **Fig. 1B-E**

**Table 1:** Comparison of conserved residues among human insulin, human IGF-1 and dcVILPs. Percentage of amino-acid residues that GIV, SGIV and LCDV-1 dcVILPs share with human insulin and IGF-1 is shown in upper panel. Percentage of important amino-acids that are important for receptor binding in dcVILPs is shown in lower panel.

| A-chain/domain |  |  |  | B-chain/domain |  |  |  |
| --- | --- | --- | --- | --- | --- | --- | --- |
|  | dc GIV | dc SGIV | dc LCDV-1 |  | dc GIV | dc SGIV | dc LCDV-1 |
| Insulin | 38% | 38% | 52% | Insulin | 30% | 33% | 47% |
| IGF-1 | 38% | 38% | 43% | IGF-1 | 45% | 45% | 59% |

| Site 1 binding residues |  |  |  | Site 2 binding residues |  |  |  |
| --- | --- | --- | --- | --- | --- | --- | --- |
|  | dc GIV | dc SGIV | dc LCDV-1 |  | dc GIV | dc SGIV | dc LCDV-1 |
| IR | 71% | 71% | 59% | IR | 43% | 43% | 64% |
| IGF1R | 52% | 52% | 48% | IGF1R | - | - | - |

### GIV and SGIV dcVILPs bind to the human insulin and IGF-1 receptors

To determine the relative affinity of dcVILPs for the two isoforms of human insulin receptor (IR-A and IR-B) and the IGF-1 receptor, we tested their ability to compete with ^125^I-Insulin and ^125^I-IGF-1 in a binding competition assay^30,31^. We used IM-9 lymphoblasts for IR-A binding competition since these cells exclusively express IR-A on their surface^32,33^. Murine embryonic fibroblasts cells derived from IGF1R knock-out mice^34^ stably transfected with either human IR-B or human IGF1R were used to assess binding competition for IR-B and IGF1R^35,36^. Consistent with previous studies^37^, we find that human IGF-1 binds to IR-A and IR-B with ~200x and ~300x lower affinity than human insulin, respectively. GIV dcVILP competes for binding to IR-A with an affinity ~3x lower than human IGF-1, while the affinity of SGIV dcVILP was comparable to IGF-1. The relative affinity of the GIV and SGIV dcVILPs for IR-B was slightly lower, with ~7-8x lower for both dcVILPs compared to IGF-1. Although LCDV-1 dcVILP has more identical residues to insulin than GIV and SGIV dcVILPs, we did not observe any binding competition with this ligand (**Fig 2A** and **B**, **Table 2**). The affinity of insulin for IGF1R was ~1000x lower compared to IGF-1, consistent with previous studies^37^. Thus, even as double chain peptides, GIV and SGIV dcVILPs had higher affinity for IGF1R than insulin by 7- to 10-fold. We did not observe any binding competition for LCDV-1 dcVILP for IGF1R (**Fig 2C**, **Table 2**).

**Figure 2:**
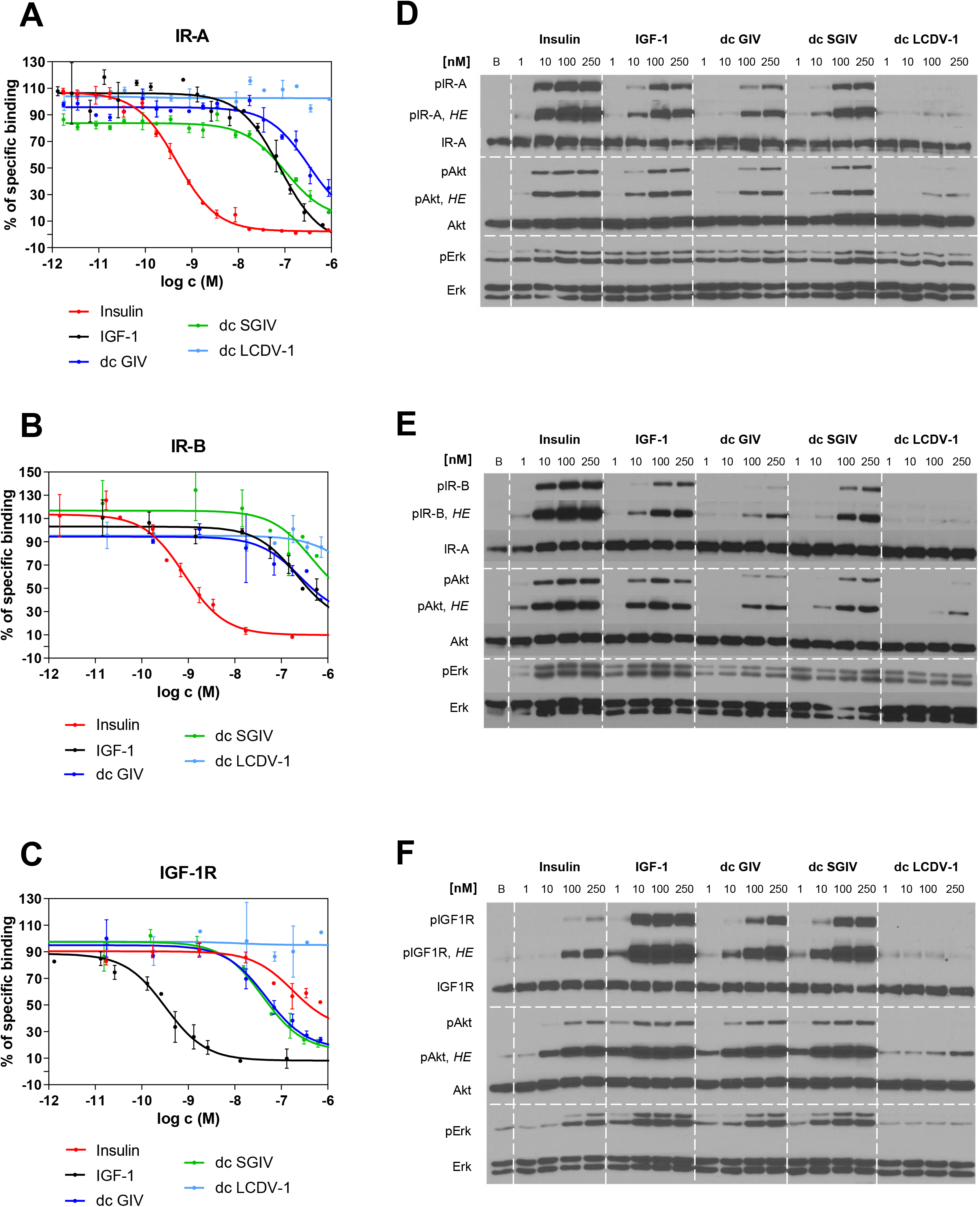
dcVILPs bind to human IR-A, IR-B and IGF1R and stimulate insulin signaling. **A - C:** Binding competition dose response curves. The curves are showing the ability of dcVILPs to compete with ^125^-I labeled human insulin for binding to IR-A **(A)** and IR-B **(B)** and with ^125^-I labeled human IGF-1 for binding to IGF1R **(C)**. IM-9 cells were used for measurements on IR-A, while murine embryonic fibroblasts derived from IGF-1 knock-out mice and stably transfected with either human IR-B or human IGF1R were used for measurements on these receptors. A representative curve for each peptide to each receptor is shown. Each point represents the mean ± SEM of duplicates. Every experiment was repeated at least three times. **D - F:** Insulin signaling via IR-A **(D)**, IR-B **(E)** and IGF1R **(F**). Murine embryonic fibroblasts derived from IGF-1 knock-out mice and stably transfected with either human IR-A, IR-B or human IGF1R were used for the experiment. Phosphorylation of the specific receptor, Akt and Erk1/2 was observed in 15 minutes after stimulation. Exposure times were between 30s to 1 min. High exposure time (*HE*) was 5 min.

**Table 2:** Receptor binding affinities of human insulin, human IGF-1 and dcVILPs to human IR-A, IR-B and IGF1R receptors. Binding affinity is reported by the equilibrium dissociation constant (K_d_). The K_d_ values were obtained from at least three independent measurements (indicated as n). Relative binding affinity is defined as K_d_ of human insulin or IGF-1/K_d_ of ligand of interest.

| Ligand | IR-A |  |  | IR-B |  |  | IGF1R |  |  |
| --- | --- | --- | --- | --- | --- | --- | --- | --- | --- |
|  | K<sub>d</sub> [nM]<br>± S.D.<br>(n) | Relative to<br>insulin<br>(fold) | Relative to<br>IGF-1<br>(fold) | K<sub>d</sub> [nM]<br>± S.D.<br>(n) | Relative<br>to<br>insulin<br>(fold) | Relative to<br>IGF-1<br>(fold) | K<sub>d</sub> [nM]<br>± S.D.<br>(n) | Relative to<br>IGF-1<br>(fold) | Relative to<br>insulin<br>(fold) |
| **Human insulin** | 0.52 ± 0.03<br>(5) | **1** | 202 | 0.58 ± 0.07<br>(3) | **1** | 303 | 293 ± 101<br>(3) | 0.0008 | **1** |
| **Human IGF-1** | 105 ± 19<br>(4) | 0.005 | **1** | 176 ± 22<br>(3) | 0.003 | **1** | 0.24 ± 0.13<br>(3) | **1** | 1221 |
| **dc GIV** | 315 ± 81<br>(3) | 0.002 | 0.33 | 1179 ± 625<br>(3) | 0.0005 | 0.15 | 41.3 ± 11.6<br>(3) | 0.006 | 7.09 |
| **dc SGIV** | 102 ± 29<br>(3) | 0.005 | 1.02 | 1441 ± 558<br>(3) | 0.0004 | 0.12 | 28.6 ± 10.0<br>(3) | 0.008 | 10.2 |
| **dc LCDV-1** | no binding<br>at 10<sup>-6</sup> M<br>(3) | - | - | K<sub>d</sub> > 10<sup>-5</sup> M<br>(3) | - | - | no binding<br>at 10<sup>-6</sup> M<br>(3) | - | - |

### dcVILPs stimulate downstream insulin/IGF-1 signaling via human IR-A, IR-B and IGF1R

To explore the effects of the dcVILPs on post-receptor signaling, we used the murine embryonic fibroblasts defined above that overexpress either human IR-A, IR-B or IGF1R^35,36^. Insulin/IGF-1 acting through their respective receptors activate (i) the PI3K/Akt pathway, that mainly regulates metabolic effects, and (ii) the Ras/MAPK pathway, that is responsible for mitogenic effects^38,39^. We tested receptor phosphorylation and phosphorylation of Akt for PI3K/Akt pathway activation and Erk1/2 for Ras/MAPK pathway activation.

On both IR isoforms, insulin induced the strongest dose-response for stimulation of the receptor autophosphorylation as expected. IGF-1 was less potent such that stimulation with 250 nM ligand was weaker than insulin at 10 nM (**Fig. 2D** **and** **E**. GIV and SGIV dcVILPs stimulated insulin/IGF signaling in a dose-dependent manner. On IR-A, SGIV dcVILP stimulated receptor phosphorylation comparable to IGF-1, and GIV dcVILP was slightly less potent at all concentrations tested (**Fig. 2D**). On IR-B, both peptides were slightly less potent compared to IGF-1 and comparable to each other (**Fig. 2E**). Both GIV and SGIV dcVILP stimulated phosphorylation of Akt and Erk1/2 in proportion to their effects on the receptor with greater effects on IR-A than IR-B (**Fig. 2D** and **2E**). Although we did not observe any competition for binding with LCDV-1 dcVILP, we observed a weak Akt and receptor autophosphorylation on both IR-A and IR-B **(Fig. 2F**). Consistent with the binding competition results on IGF1R, SGIV and GIV dcVILPs stimulated post-receptor signaling more potent than insulin, with SGIV dcVILP being slightly more potent than GIV dcVILP. As observed for the IR, although we did not observe any binding competition for LCDV-1 dcVILP, it stimulated a weak signal for receptor and Akt phosphorylation (**Fig. 2F**).

### GIV and SGIV dcVILPs are active in vivo and stimulate glucose uptake in mice

To test whether dcVILPs can stimulate glucose uptake in vivo, we performed an insulin tolerance test (ITT). Adult C57BL/6J mice were injected intraperitoneally with either 6 nmol/kg insulin or different concentrations of GIV and SGIV dcVILPs (**Fig. 3**). Based on our in vitro data showing about 0.05 % relative affinity for the IR, we decided to use 0.3 μmol/kg (50x higher concentration than insulin) GIV and SGIV dcVILPs in male mice. Consistent with previous studies^9,40^, insulin caused ~ 60% decrease in blood glucose in 60 minutes, after which glucose started to increase. Surprisingly, injection with 50x GIV or SGIV dcVILPs led to very severe hypoglycemia such that we needed to terminate the experiment in 30 minutes by injecting glucose to save the animals (**Fig. 3A**).

**Figure 3:**
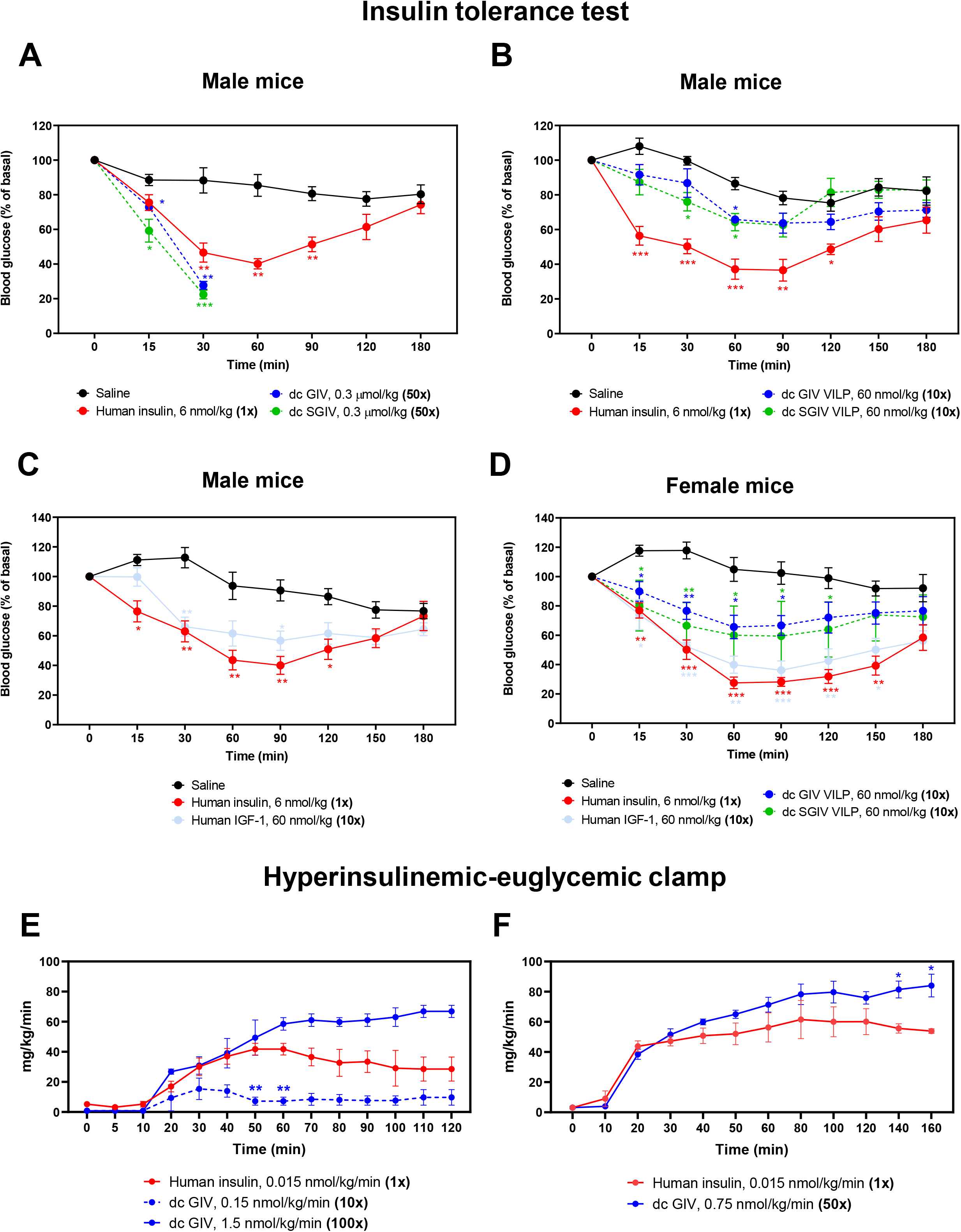
GIV and SGIV dcVILPs stimulate glucose uptake in mice. **A – D:** Insulin tolerance test. C57BL/6J mice were injected i.p with human insulin, human IGF-1, GIV and SGIV dcVILPs or saline. The concentration of insulin was 6 nmol/kg in all panels, whereas the concentration of dcVILPs was 0.3 μmol/kg **(A**) and 60 nmol/kg **(B, D).** Concentration of human IGF-1 was 60 nmol/kg **(C, D).** Blood glucose was measured within the range from 0 to 180 minutes. Data are mean ± S.E.M. (*P<0.05; **P<0.01, *** P < 0.001). Mixed-effects analysis followed by Dunnett’s multiple comparisons test was applied, n = 5 in all groups. **E:** Glucose infusion rates during the 2-hour in vivo experiments with infusion of human insulin or GIV dcVILP. The concentration of insulin was 0.015 nmol/kg/min, and the concentration of GIV dcVILP was 0.15 and 1.5 nmol/kg/min. n = 4 for insulin and n = 2 for both concentrations of GIV dcVILP. **F:** Glucose infusion rates during the 3-hour in vivo experiments with infusion of human insulin or GIV dcVILP. The concentration of insulin was 0.015 nmol/kg/min, and the concentration of dc GIV dcVILP was 0.75 nmol//kg/min. n = 4 for both groups. Two-way repeated measures ANOVA followed by Tukey’s multiple comparisons test was applied. Data are mean ± S.E.M (*P<0.05; **P<0.01).

When this experiment was repeated using 60 nmol/kg concentrations of the dcVILPs (10x) (**Fig. 3B** and **D**), again, both dcVILPs were able to significantly lower the blood glucose, and at 60 minutes produced 57-58% of the effect of insulin (**Fig. 3B**). By comparison, injection of IGF-1 at 60 nmol/kg concentration (10x) reached 71% of the effect of insulin at 60 min, and this effect persisted for longer than insulin’s effect consistent with the longer half-life of IGF-1 compared to insulin (**Fig. 3C**). Similar results were obtained in female mice (**Fig. 3D**). Taken together, these results indicate that both dcVILPs have in vivo glucose lowering effects similar to IGF-1, but slightly more than an order of magnitude less potent than insulin and with a longer duration of effect.

### In vivo infusion experiments reveal white adipose tissue specific effects of GIV dcVILP

To further explore the mechanism of dcVILP action in vivo, we performed an in vivo experiment using an acute administration of GIV dcVILP in awake mice. In this experiment, male C57BL/6J mice were intravenously infused with constant levels of tested ligand or insulin for 2 hours, and 20% glucose was infused at variable rates to maintain euglycemia. We first optimized the conditions using a 0.015nmol/kg/min dose for insulin and 0.15 nmol/kg/min (10x) and 1.5 nmol/kg/min (100x) for GIV dcVILP. GIV dcVILP at 100x induced a strong glucose disposal as reflected by a profound increase in glucose infusion rate during the experiments, and there was a dose-dependent effect of GIV dcVLIP as shown by a minimal effect of GIV dcVILP at 10x on glucose disposal in mice (**Fig 3E**). Based on this dose optimization experiment, we performed a 3-hour infusion of 0.75 nmol/kg/min (50x) concentration of the GIV dcVILP and compared the effects to a 3-hour infusion of insulin at 0.015 nmol/kg/min in awake mice. Our data indicate that GIV dcVILP at 50x induced an increase in glucose disposal similar to insulin, as reflected by comparable rates of glucose infusion during the 3-hour experiments (**Fig. 3F**).

In additional cohort of mice, we performed a 3-hour infusion of 0.75 nmol//kg/min (50x) concentration of the GIV dcVILP or insulin at 0.015 nmol/kg/min with a continuous infusion of [3-^3^H]glucose to assess whole body glucose turnover, and 2-deoxy-D-[1-^14^C]glucose was administered as a bolus at 45 min before the end of experiments to measure glucose uptake in individual organs. Measurements of glucose uptake in heart, skeletal muscle (gastrocnemius), brown adipose tissue (BAT, intrascapular) and white adipose tissue (WAT, epididymal) identified a unique characteristic of GIV dcVILP. While GIV dcVILP (50x) stimulated a comparable glucose uptake compared to insulin in gastrocnemius muscle, heart and BAT (**Fig. 4A-C**)., the glucose uptake was significantly (1.9 fold) increased in WAT compared to insulin (**Fig. 4D**). This finding suggests a tissue selective effect for GIV dcVILP on glucose metabolism in white adipose tissue.

**Figure 4:**
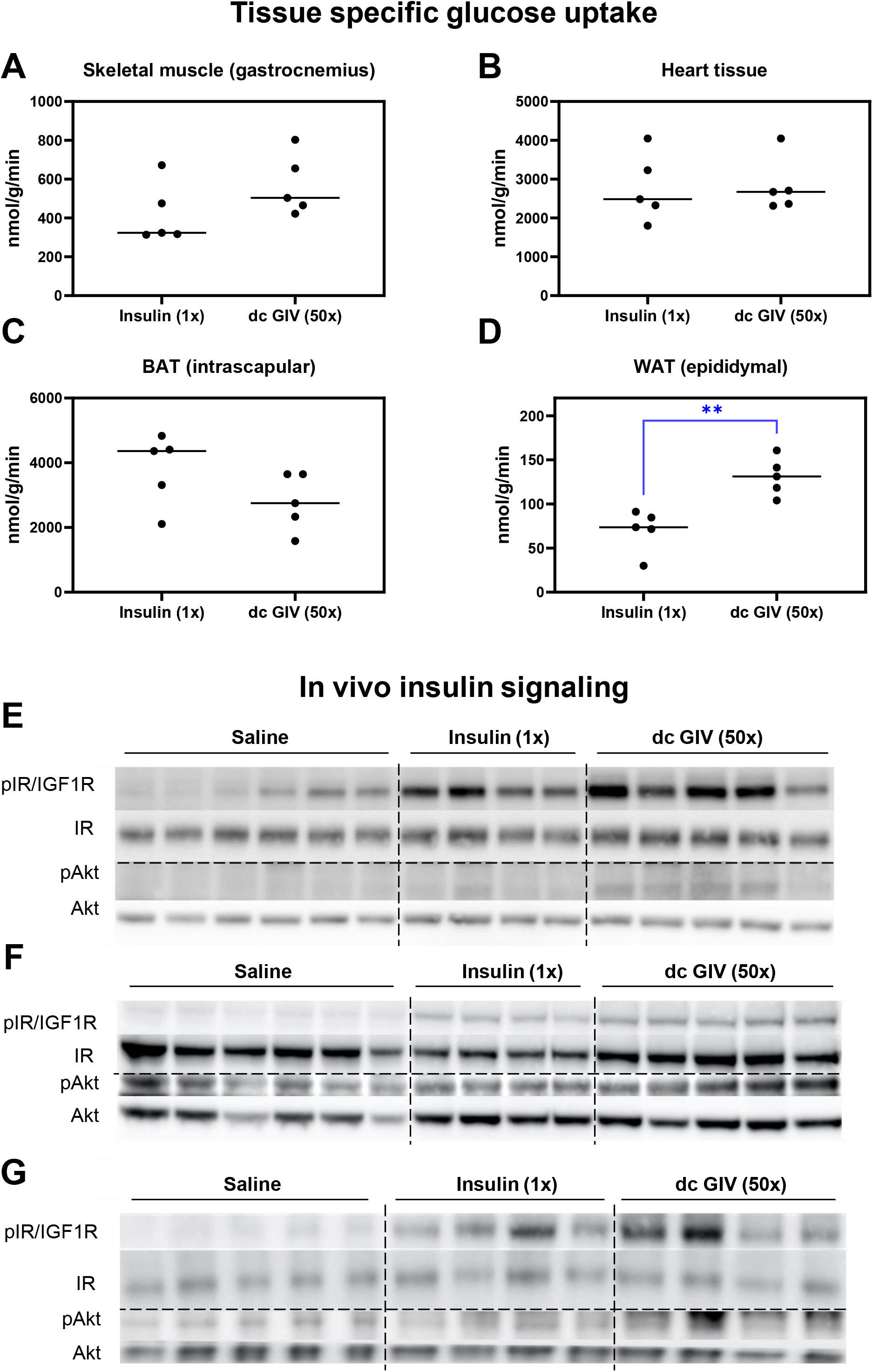
GIV dcVILP stimulates in vivo insulin signaling and WAT specific glucose uptake. **A** - **D:** Tissue-specific glucose uptake after a 3-hour infusion of insulin (0.015 nmol/kg/min; 1x) or GIV dcVILP (0.75 nmol//kg/min; 50x) in awake mice (n=5 for each group). Student’s t-test was applied (**P<0.01). n=5 for both groups. **E – G:** in vivo signaling in insulin sensitive tissues obtained at the end of 3-hour infusion of insulin (0.015 nmol/kg/min; 1x) or GIV dcVILP (0.75 nmol//kg/min; 50x) in awake mice. Basal tissue samples were collected after a 3-hour saline infusion. **E:** liver, **F:** skeletal muscle (gastrocnemius), G: WAT (epididymal).

Hepatic glucose production was significantly suppressed in both insulin and GIV dcVILP groups but we did not determine any significant difference related to insulin action in the liver (**Table S1**). In separate experiments, we assessed insulin signaling in liver, gastrocnemius muscle and WAT and found that 50x GIV dcVILP stimulated phosphorylation of IR/IGF1R and Akt in all tissues (**Fig. 4E-G**, **Fig. S2**). Akt phosphorylation was significantly increased by GIV dcVILP in the liver (p=0.0036) and WAT (p=0.0009) compared to the insulin group (**Fig. S2B, F**).

### GIV dcVILP induces GLUT4 gene expression in WAT in vivo

To further understand the tissue selective effects of the GIV dcVILP observed for WAT, we used tissues collected at the end of a 3-hour in vivo infusion of GIV VLIP or insulin in awake mice to evaluate the expression of the receptors and insulin-stimulated genes. Basal tissue samples were collected from mice after a 3-hour infusion of saline. Because our in vitro data showed that GIV dcVILP stimulates IGF1R more than insulin, we first explored the possibility that the GIV dcVILP specific glucose uptake is caused by different receptor composition in different tissues. Using RT-qPCR, we showed that liver, BAT and WAT contain the highest amount of IR-B and low amounts of IR-A and IGF1R. In contrast, the most abundant receptor in skeletal muscle was IR-A (**Fig. S3**). These results showed that higher glucose uptake is not related to increased IGF1R expression in WAT. Next, we evaluated the expression of genes related to insulin action in liver, skeletal muscle (quadriceps), BAT and WAT. Specifically, we focused on the genes related to glucose metabolism and lipogenesis. We also tested GLUT4 in all four tissues, and thermogenesis marker uncoupling protein 1 (UCP-1) in BAT.

Consistent with our findings from in vivo glucose uptake, GLUT4 expression was significantly higher (1.5 fold) in GIV dcVILP-stimulated WAT compared to insulin-stimulated WAT (**Fig. 5A**). In addition to GLUT4, fatty acid synthase (FASN) expression was increased by 2.1 fold in GIV dcVILP compared to insulin (**Fig. 5B**). Insulin and GIV dcVILP stimulated sterol regulatory element-binding protein 1-C (SREBP-1c) expression in a similar manner (**Fig. 5C**), while GIV dcVILP showed an increasing trend for acetyl-CoA carboxylase 1 (ACACA) expression compared to insulin (p=0.087, **Fig. 5D**). We did not observe any significant differences between insulin and GIV dcVILP stimulated gene expression in any of the genes tested in muscle (**Fig. S4**) and BAT (**Fig. S5**).

**Figure 5:**
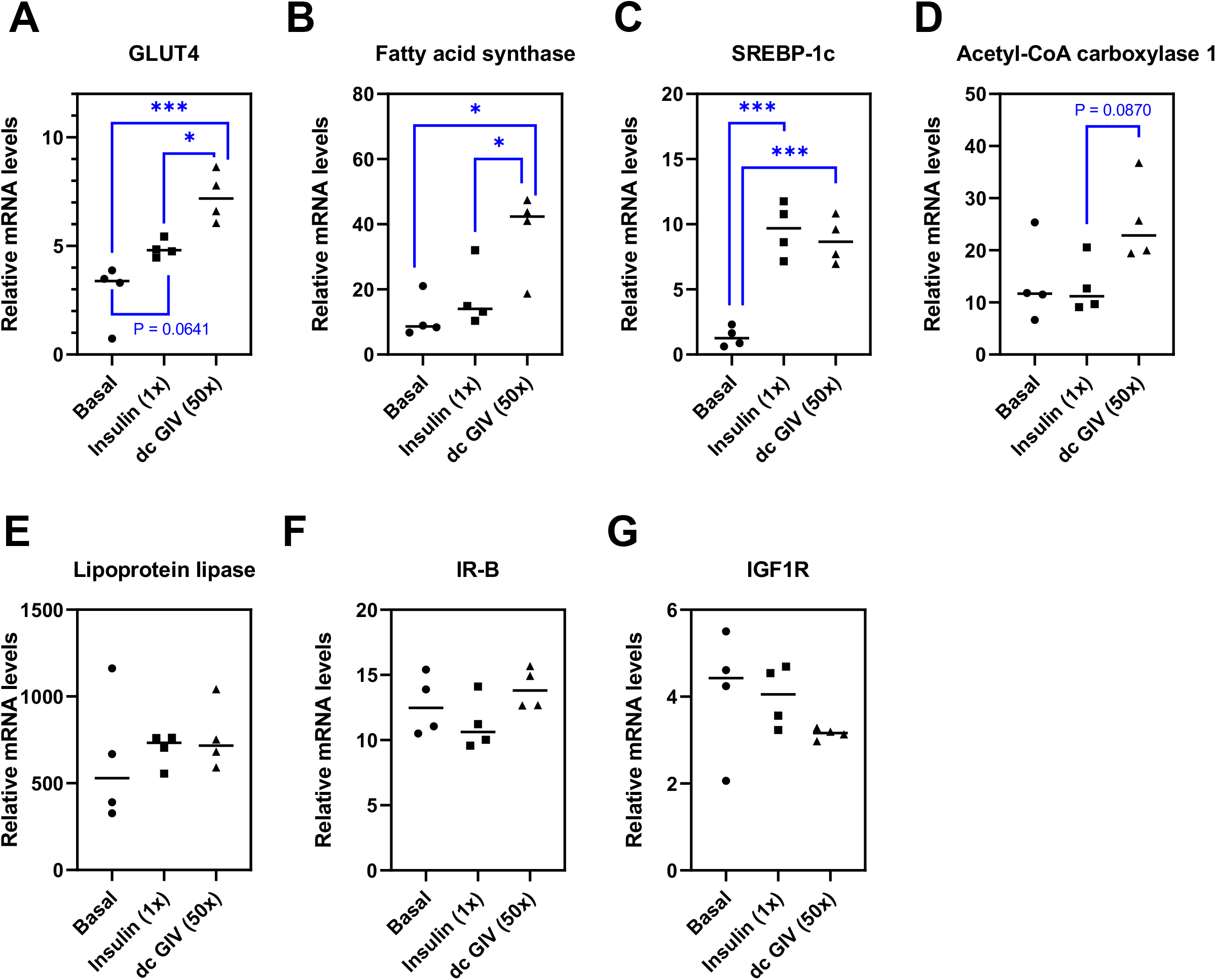
RT-qPCR analysis of expression of genes connected with insulin function and lipogenesis in murine WAT. Tissues were collected after 3-hour insulin (0.015 nmol/kg/min; 1x) or GIV VILP (0.75 nmol/kg/min; 50x) infusion, and basal tissue samples were collected after 3 hours of saline infusion in awake mice. Data are expressed as % of β-actin. n = 4 per group. Ordinary one-way ANOVA followed by Tukey’s multiple comparison test was applied (*P<0.05; **P<0.01, ***P<0.001). P-values lower than 0.1 are indicated.

Consistent with previous studies on insulin action in the liver^41^, the gluconeogenesis markers, catalytic subunit of glucose-6-phospgatase (G6PC) and phosphoenol pyruvate carboxykinase 1 (PCK1), were downregulated by both insulin and GIV dcVILP (**Fig. 6A** and **B**), while glucokinase (GCK), the glycolysis marker, was upregulated (**Fig. 6C**). Interestingly, we observed a significant (1.6 fold) increase for GCK in the GIV dcVILP group compared to the insulin group. The lipogenesis markers were decreased by both insulin and GIV dcVILP (**Fig. 6D-G**). Although, the GLUT4 expression is very low in the liver^42^, we observed a significant increase in GLUT4 expression after stimulation by GIV dcVILP when compared to both saline as well as insulin (4.7 fold) (**Fig. 6H**). When we analyzed the receptor expression, both insulin and GIV dcVILP downregulated IR-B expression in the liver (**Fig. 6I**), whereas only GIV dcVILP downregulated IGF1R expression (**Fig. 6J**).

**Figure 6:**
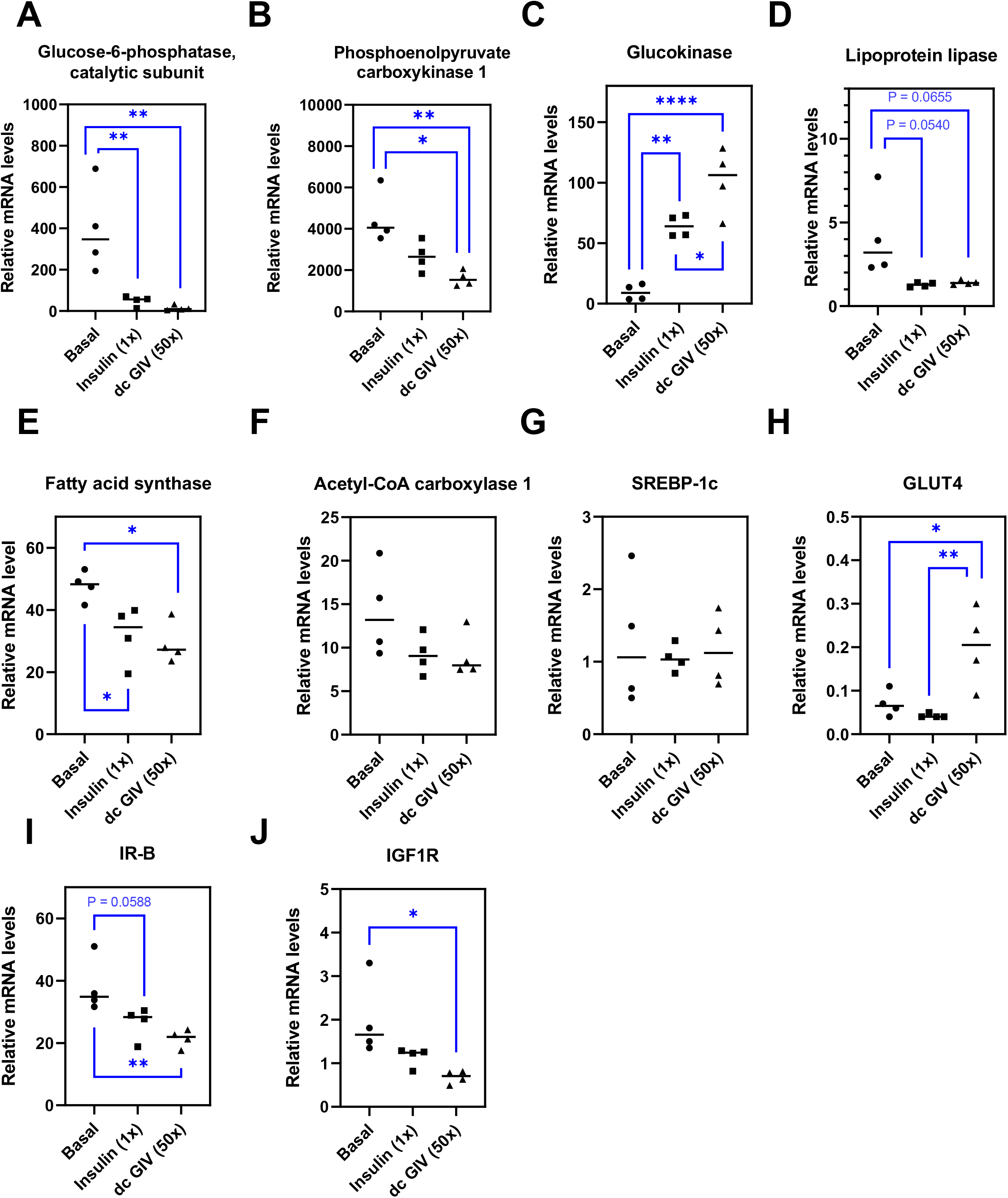
RT-qPCR analysis of expression of genes connected with insulin function, and lipogenesis in murine liver. Tissues were collected after 3-hour insulin (0.015 nmol/kg/min; 1x) or GIV VILP (0.75 nmol/kg/min; 50x) infusion, and basal tissue samples were collected after 3 hours of saline infusion in awake mice. Data are expressed as % of β-actin. n = 4 per group. Ordinary one-way ANOVA followed by Tukey’s multiple comparison test was applied (*P<0.05; **P<0.01, ***P<0.001, ****P<0.001). P-values lower than 0.1 are indicated.

## DISCUSSION

We recently discovered that four viruses belonging to the *Iridoviridae* family possess genes with high homology to human insulin and IGFs^9^. In this study, we characterized three of these VILPs in their insulin-like, i.e. double-chain, forms for the first time. We first showed that GIV and SGIV dcVILPs can bind to IR-A, IR-B and IGF1R. Interestingly, on IR-A, the affinity of SGIV dcVILP is comparable to that of IGF-1, whereas the affinity of GIV dcVILP is ~ 3x lower. On IR-B, however, the affinities of GIV and SGIV dcVILPs are comparable to each other and ~ 7-8x lower than IGF-1. The only difference between IR-A and IR-B is 12 extra amino acids in the α-CT peptide that are present in IR-B but not IR-A^32,35^. This α-CT peptide is directly involved in ligand binding^21,22,24,25^.

The sequences of GIV and SGIV dcVILPs differ only in three amino acids (**Fig. S1**), which correspond to insulin residues ValB2, ProB28 and SerA12. ValB2 and ProB28 have not been shown to be involved in the insulin:IR interaction, whereas SerA12 was shown to be involved in the Site 2 interaction^24,25^. SerA12 substitution by alanine decreases the affinity to IR to 36% of insulin^27^, however, SerA12 is known to be involved in interaction within IR FnIII-1 domain^24,25^. Therefore, SerA12 is unlikely to play a role in the differential binding to IR-A and IR-B. The only residue that lies in a region that is involved in interaction with the α-CT peptide (specifically C-terminal region of insulin B-chain^21,24,25^) is the residue corresponding to insulin ProB28. This residue is substituted by serine in GIV dcVILP and proline in SGIV dcVILP. Therefore, it is probable that the ProB28Ser substitution lies behind the decreased affinity of GIV dcVILP to IR-A compared to SGIV dcVILP. By modeling of GIV and SGIV dcVILPs onto Site 1 of IR, we showed that the ProB28 presence makes the following ArgB28 and ArgB29 direct to the α-CT helix segment in SGIV dcVILP, while these residues are directed away from it in GIV dcVILP (**Fig. S6A**). Importantly, the last receptor amino acid in the model is Arg717 (PDB 6PVX) which is the last amino acid that is identical in IR-A and IR-B. After Arg717, there are three additional amino acids in IR-A, while there are 15 more amino acids in IR-B that are not present in the model. Therefore, GIV dcVILP would potentially clash with the following α-CT sequence in both IR-A and IR-B, while SGIV dcVILP would avoid this clash in IR-A, but would still clash with the longer IR-B. This may explain why SGIV dcVILP has three-fold higher affinity for IR-A than GIV dcVILP, while their affinity for IR-B is comparable.

Another interesting observation is that GIV and SGIV dcVILPs have higher affinity to bind and stimulate signaling via IGF1R than insulin, since these ligands are missing the C-domain that is important for IGF-1:IGF1R interaction^43–46^. Even though dcVILPs are completely missing the C-domain, they bind to IGF1R with 7x (GIV dcVILP) and 10x (SGIV dcVILP) higher affinity compared to insulin. Signaling experiments are consistent with the binding results and showed a similar trend as both GIV and SGIV dcVILPs stimulated phosphorylation of IGF1R, Akt and Erk with higher potency than insulin. The comparison of amino acid sequences of GIV and SGIV dcVILPs with insulin and IGF-1 revealed that several amino-acids that are involved in insulin binding to the Site 2 of IR are replaced in these VILPs by amino acids that are identical to IGF-1 in the corresponding positions - and differ from insulin. Specifically, these include GluB10 (corresponding to Glu9 in IGF-1 and HisB10 in insulin), AspB13 (corresponding to Asp12 in IGF-1 and GluB13 in insulin) and AspB21 (corresponding to Asp20 in IGF-1 and GluB21 in insulin) (**Fig 1A**). Interestingly, two of these three residues in IGF-1 (Glu9 and Asp12) are involved in IGF1R Site 1 binding. Comparisons of these two residues in models of GIV bound to the Site 2 of IR and Site 1 of IGF1R to human insulin (**Fig. S6B)** and IGF-1 (**Fig. S6C**) indicate that GIV and SGIV dcVILPs might preferentially bind to the Site 1 of IGF1R than to Site 2 of IR. This may be a possible explanation of why GIV and SGIV dcVILPs show an increased affinity and ability to activate IGF1R than insulin. Moreover, the HisB10Glu/Asp mutation in insulin is itself well known for dramatically enhancing the binding affinity to both IGF1R and IR-A^47–49^.

One of the most interesting findings of this study is related to analysis of glucose uptake in the in vivo infusion experiments. We showed that GIV dcVILP (50x) specifically stimulates ~2 fold glucose uptake in epididymal white adipose tissue compared to insulin. We first explored the distribution of the receptors in insulin sensitive tissues, but we did not observe an increased expression of IGF1R in WAT. Because we determined an increased Akt phosphorylation for GIV dcVILP compared to insulin in WAT, we decided to investigate the genes related to insulin action. We observed an increased GLUT4 and FASN expression in the GIV dcVILP group compared to insulin. This was specific to WAT and not observed in BAT and skeletal muscle. In liver, glucokinase was significantly increased in mice receiving GIV dcVILP compared to insulin. The increase in glucokinase might be related with using an increased dose of GIV dcVILP compared to insulin.

Our results on GLUT4 expression are particularly interesting. Since its discovery in 1988^50^, there have been tremendous efforts to understand the function and regulation of GLUT4^51^. Insulin is known to stimulate GLUT4 translocation^51,52^, but not GLUT4 expression. Our data show that GIV dcVILP significantly stimulates GLUT4 expression in WAT. Further, both insulin and GIV dcVILP stimulated GLUT4 expression in BAT. These results indicate that GLUT4 expression in adipose tissue can be regulated by specific insulin analogues. It is previously shown tht GLUT4 expression is decreased in obesity and increased in response to exercise adipocytes^53^. Further, overexpression of GLUT4 in adipose tissue makes mice more insulin sensitive and glucose tolerant^54,55^. Thus, identification and synthesis of novel insulin analogues targeting GLUT4 expression in adipose tissue might be a novel approach to be tested in diabetes control in the future.

According to our knowledge, GIV dcVILP is the first insulin analogue that has WAT specificity and further studies are needed to explain the specific mechanism underlying the tissue-selectivity. Previous studies have identified hepatoselective action for different insulin analogues^56–61^. This selectivity is thought to be related with either increased molecular size (proinsulin and insulin peglispro) or their ability to bind endogenous proteins (thyroxyl conjugates and insulin detemir)^62^. The increased data produced by genome projects have increased our ability to understand the natural repertoire of hormone ligands. For example, the Gila monster exendin-4 mimics GLP-1 functions and unlike human GLP-1, it has a long half time^63,64^. Likewise, recent discovery of cone snail venom insulins have potential to help us designing fast-acting insulin analogues^65–67^. Thus, characterization of new VILPs that are evolved as a result of host-pathogen interactions, and understanding the characteristics of tissue specificity, has potential to help us design better insulin therapies.

In our previous study, we showed that the sequences of these VILP-carrying viruses are identified in human fecal and plasma samples^9^. Although this finding suggests that humans are exposed to these viruses, it is still unclear whether these fish viruses can infect humans. While we continue to work on this question, if they do infect humans, this will raise several questions regarding their link to human disease including diabetes, cancer and hypoglycemia. While the number of viruses that can infect mammalian animals are predicted to be over 320,000^68^, there are only 10,316 complete viral genomes in the NCBI database as of August 1, 2020. Thus, we expect to identify human viruses carrying VILPs in the future. Human viruses are known to target cellular metabolism by changing the expression levels of transcription factors, metabolic intermediates and enzymatic activity^69–72^. We anticipate that VILP carrying viruses are targeting the glucose metabolism and cell cycle when they infect fish to promote their replication. Furthermore, insulin and IGF-1 are mitogenic and anti-apoptotic molecules^73^ that are two perfect characteristics that a pathogen needs. Indeed, overexpression of SGIV VILP stimulated cell proliferation in fish cells and increased SGIV replication^74^.

Taken together, our study shows that GIV and SGIV dcVILPs are new members of the insulin/IGF superfamily with remarkable in vitro and in vivo insulin/IGF-1 like effects. Although we could not show binding competition for LCDV-1 dcVILP to human IR and IGF1R, it can stimulate a weak signal that needs further investigation. Identification of tissue selectivity of GIV dcVILP has potential to help us to better understand the tissue selectivity of insulin. Furthermore, the effects of GIV dcVILP on GLUT4 expression opens a new avenue to better understanding of Glut4 regulation by insulin action. In summary, our findings contribute to our understanding of VILP action on human receptors and have potential to help with designing new insulin/IGF analogs specific to WAT.

## MATHERIALS AND METHODS

### Bioinformatics

The sequence alignments presented in this paper were prepared using a multiple sequence alignment program (Clustal Omega). We used the website https://swissmodel.expasy.org/ for the homologous building^75^, and to align the modelled dcVILPs with insulin or IGF-1 in IR (PDB: 6PXV) or IGF1R structures (PDB: 6PYH). The final figures prepared using PyMOL.

### Peptide synthesis

Viral insulin-like peptides were synthesized via Fmoc solid phase peptide synthesis (SPPS) utilizing a commercial automated peptide synthesizer (Symphony® X, Gyros Protein Technologies) using a similar method to what has been previously reported^76^. Briefly, A- and B-chains were individually synthesized at a 0.1 mmol scale using standard Fmoc protected amino acids, a specific set of orthogonally side-chain protected cysteine residues that allows for directed disulfide bridge formation, pseudoproline and isoacyl dipeptide building blocks that aid in overcoming coupling difficulties during SPPS and ameliorate solubility issues during HPLC purification, respectively. Fmoc deprotection was carried out using 20% Piperidine in DMF, and coupling reactions were done using DIC/Oxyma for 1 hour using a 9-fold excess of reagents. Fmoc-Rink-MBHA resin was used as the solid support for the synthesis of the A-chains. The Cysteine protection arrangement used for GIV and SGIV dcVILPs A-chains was as follows: Cys(StBu)A6, Cys(Acm)A7, Cys(Mmt)A11, and Cys(Trt)A20. The Cysteine protection arrangement used for LCDV-1 dcVILP A-chain was as follows: Cys(STmp)A6, Cys(Acm)A7, Cys(Mmt)A12, and Cys(Trt)A21. While no isoacyl dipeptides were used for GIV and SGIV dcVILPs A-chains, the LCDV-1 dcVILP A-chain required the use of Boc-Thr(Fmoc-Ala)-OH isoacyl dipeptide at positions A3,4 and Fmoc-Thr(tBu)-Thr(^ψMe,Me^Pro)-OH pseudoproline dipeptide at positions A8,9 and A13,14. A-chains are synthesized bearing the intramolecular disulfide bridge (CysA6-CysA11 for GIV and SGIV dcVILPs and CysA6-CysA12 for LCDV-1). Said bond was formed during the SPPS process as outlined in scheme 2 as reported by Liu et al.^76^. A slight deviation from scheme 2 was necessary for the synthesis of LCDV-1 dcVILP A-chain in that Cys(STmp)A6 was deprotected with 0.1 M N-methylmorpholine and 5% dithiothreitol in DMF instead of 25% β-mercaptoethanol in DMF. B-chains for GIV and SGIV dcVILPs were synthesized on Fmoc-Arg(Pbf)-Wang resin and the B-chain for LCDV-1 dcVILP was carried out using H-Thr(tBu)-HMPB-ChemMatrix resin. The Cysteine protection arrangement used for GIV, SGIV and LCDV-1 dcVILPs B-chains was as follows: Cys(Acm)B7, and Cys(Trt)B19. While no isoacyl dipeptide was used for the LCDV-1 dcVILP B-chain, GIV and SGIV dcVILPs B-Chains included the use of Boc-Thr[Fmoc-Tyr(tBu)]-OH isoacyl dipeptide at positions B25,26. The final solid support cleavage and global side chain deprotection is achieved using standard trifluoroacetic acid (TFA) mediated acidolysis protocols, with the inclusion of DTNP in the cleavage cocktail of the B-chains to afford the activated Cys(SNpy)19 residue. The A- and B-chains were purified using standard reverse phase HPLC methods (TFA acidified water/acetonitrile mobile phases) and were lyophilized to dryness after purification. The intermolecular disulfide bridges CysA20-CysB19, ACys7-CysB7 for GIV and SGIV dcVILPs; and CysA21-CysB19, CysA7-CysB7 for LCDV-1 dcVILPs were formed in a guided and sequential manner exploiting the orthogonality of the cysteine protection scheme. A-chains, B-chains, intermediates, as well as the final dcVILPs were characterized by analytical LC-MS, purified by RP-HPLC and lyophilized to dryness.

### Cell culture

Human IM-9 lymphocytes (ATCC) and murine embryonic fibroblasts, that were derived from IGF1R knockout mice and stably transfected with either IR-A (R^−^/IR-A cells), IR-B (R^−^/IR-B cells) or IGF1R (R^+39^ cells), kindly provided by A. Belfiore (Catanzarro, Italy) and R. Baserga (Philadelphia, PA), were cultured as described previously^30,77^.

### Receptor binding studies

For receptor binding studies, human IM-9 lymphoblasts, that express IR-A exclusively, and R^−^/IR-B and R^+39^ murine embryonic fibroblasts (described above) were used for a whole-cell receptor-binding assay. Receptor binding assays with IR-A were perfomed accoding to Morcavallo et al.^30^ and binding assays with IR-B and IGF1R were performed according to Kosinova et al.^31^. The binding curve of each ligand was determined in duplicate, and the final dissociation constant (K_d_) was calculated from at least three (n ≤ 3) binding curves. Human insulin and human IGF-1 were supplied by Merck. Human ^125^I-insulin (NEX420050UC) and human ^125^I-IGF-1 (NEX241025UC) were supplied by Perkin-Elmer.

### Receptor phosphorylation and downstream signaling

For receptor phosphorylation and downstream signaling experiments, R^−^/IR-A, R^−^/IR-B and R^+39^ murine embryonic fibroblasts (described above) were used to explore signaling properties of ligands via specific receptors. Cells were seeded into 24-well plates (Denville Scientific) (8×10^4^ cells per well) in 300 μl of DMEM media (Corning) and grown overnight. Afterwards, cells were washed twice with PBS and starved in serum-free media for 4 hours. After the starvation, cells were washed with pure DMEM media and incubated with ligand diluted in pure DMEM media (0, 1, 10, 100 and 250 nM) in 37°C for 15 min. The reaction was terminated by washing the cells with ice-cold PBS (HyClone) followed by snap freezing in liquid nitrogene. Cell lysis was performed using 50 μl of RIPA buffer (Millipore) supplemented with protease and phosphatase inhibotors (Bimake). Cells on plates were incubated in the RIPA buffer on ice for 15 minutes, then transferred to microtubes and incubated on ice for additional 15 minutes. The lysates were centrifuged (13 000g, 5 min, 4°C) and supernatant was transferred to new microtubes. Protein concentration in each sample was evaluated using BCA Assay (Thermo Fisher Scientific). Samples were further diluted using sample buffer for SDS-PAGE (final concentration 62.5 mM Tris, 2% SDS (w/v), 10% glycerol (v/v). 0.01% bromphenol blue (w/v), 0.1M DTT (w/v), pH = 6.8 (HCl)) and routinely analyzed using SDS-PAGE and immunoblotting. Cell lysates (4 μg of protein content/sample) were separated on 10% polyacrylamide gels and electroblotted to PVDF membrane. The membranes were probed with primary antibodies against phospho-IR/IGF1R (1:500, #3024), phospho-Akt (S473) (1:1000, #9271) and phospho-Erk1/2 (T202/Y204) (1:5000, #9101). All primary antibodies against phospho-proteins were purchased from Cell Signaling Technology. The western blots were developed using SuperSignal West Pico PLUS Sensitivity substrate (Thermo Fisher Scientific). For detection of the amount of total proteins, standard stripping procedure using the mild stripping buffer (1.5% glycine (w/v), 0.1% SDS (w/v), 1% Tween20 (v/v), pH=2.2 (HCl)) was used and the membranes were relabeled with primary antibodies against IRβ (1:1000, #3025), IGF1Rβ (1:1000, #9750), Akt (1:1000, #4685) and Erk1/2 (1:2000, #9102). All primary antibodies against total proteins were purchased from Cell Signaling Technology. HRP Goat Anti-Rabbit secondary antibody was used in all cases (1: 10000, ABclonal #AS014).

### Insulin tolerance test

All animal studies presented in this study complied with the regulations and ethics guidelines of the NIH and were approved by the Boston College Institutional Animal Care and Use Committee. Insulin tolerance testing was performed on 12 to 20-week-old male C57BL/6J mice (Jackson Laboratory). Mice were grouped according to their weight before experiment. After 4-hour starvation, mice were injected i.p. with insulin (Humulin, 6 nmol/kg (corresponds to 1.0 U/kg) (Eli Lilly), GIV and SGIV dcVILPs (0.3 μmol/kg or 60 nmol/kg), LCDV-1 dcVILP (1 μmol/kg) and saline as a control (n = 5 per condition). Tail-vein blood glucose was measured at the indicated time points (**Fig. 3**) using an Infinity glucometer (US Diagnostic Inc.). Statistical analysis was done using Mixed effects analysis - Dunnett’s multiple comparisons test.

### In vivo infusion experiments in awake mice

All in vivo infusion experiments in mice were conducted at the National Mouse Metabolic Phenotyping Center (MMPC) at UMass Medical School, and animal studies were approved by the Institutional Animal Care and Use Committee of the University of Massachusetts Medical School. Male C57BL/6J mice received a survival surgery to establish an indwelling catheter in the right internal jugular vein. After recovery of 4-5 days, mice were fasted overnight (~16 hours) and placed in rat-sized restrainers for in vivo experiments. In the dose optimization experiment, mice received a continuous infusion of insulin (0.015 nmol/kg/min, corresponds to 2.5 mU/kg/min, n=4) or GIV (0.15 nmol/kg/min or 1.5 nmol/kg/min, n=2 for each group) for 2 hours, and 20% glucose was infused at variable rates to maintain euglycemia. Blood samples were collected from the tail tip at 10 min intervals to measure plasma glucose levels during the 2-hour experiments.The experiment was later repeated using continuous infusion the same concentration of insulin (0.015 nmol/kg/min, n = 4), GIV dcVILP (0.75 nmol/kg min, n = 4) or saline (n = 4).

Additional cohort of male C57BL/6J mice received a continous infusion of insulin (0.015 nmol/kg/min, n=5), GIV dcVILP (0.75 nmol/kg/min, n = 5) or saline (n=6) for 3 hours, and 20% glucose was infused at variable rates to maintain euglycemia. During the experiments, [3-^3^H]glucose (PerkinElmer, Waltham, MA) was continuously infused for 3 hours to assess whole body glucose turnover, and 2-deoxy-D-[1-^14^C]glucose was administered as a bolus (10 μCi) at 45 min before the end of experiments to measure glucose uptake in individual organs^78^. Blood samples were collected from the tail tip at 10-20 min intervals during the experiments. At the end of experiments, mice were euthanized, and tissues (skeletal muscle, liver, brown and white adipose tissue and heart) were harvested, snap frozen in liquid nitrogen and kept at −80°C for biochemical analysis. Statistical analysis was done using Two-way repeated measures ANOVA followed by Tukey’s multiple comparisons test.

### Biochemical analysis of glucose metabolism

Glucose concentrations were analyzed using 5 μl plasma by a glucose oxidase method on Analox GM9 Analyser (Analox Instruments Ltd., Hammersmith, London, UK). Plasma concentrations of [3-^3^H]glucose and 2-deoxy-D-[1-^14^C]glucose were determined following deproteinization of plasma samples as previously described^78^. For the determination of tissue 2-[^14^C]DG-6-phosphate (2-[^14^C]DG-6-P) content, tissue samples were homogenized, and the supernatants were subjected to an ion exchange column to separate 2-[^14^C]DG-6-P from 2-[^14^C]DG. Glucose uptake in individual tissues was assessed by determining the tissue content of 2-[^14^C]DG-6-P and plasma 2-[^14^C]DG profile.

### Molecular analysis using tissues collected from in vivo infusion experiments

The following molecular analysis (insulin signaling, RNA isolation, and RT-qPCR) was performed using tissue samples collected from in vivo infusion experiments at the National MMPC at UMass Medical School. Male C57BL/6J mice received a continous infusion of insulin (0.015 nmol//kg/min) or GIV dcVILP (0.75 nmol/kg/min), or saline (n = 4-6) for 3 hours, and 20% glucose was infused at variable rates to maintain euglycemia. Blood samples were collected from the tail tip at 10-20 min intervals during the experiments. Basal tissue samples were collected after a 3 hour infusion of saline in awake mice.

### In vivo insulin signaling

Tissues were lysed in RIPA buffer (EMD Millipore) supplemented with 0.1% SDS and a cocktail of protease and phosphatase inhibitors (Biotools). Proteins were denatured in denaturing buffer (NuPAGE LDS Sample Buffer, Thermo Fisher Scientific) supplemented with 5% of β-mercaptoethanol and incubated at 90°C for 5 minutes. 10 μg/well of protein was loaded on a 4-12% NuPAGE Bis-tris gel (Thermo Fisher Scientific) and then transferred on PVDF membrane (Thermo Fisher Scientific). Membrane was blocked in blocking buffer (Thermo Fisher Scientific) for 1h at room temperature and incubated with primary antibody (1:1000) over-night and with secondary antibody (1:1000) for 4 hours. The membranes were probed with the following antibodies: IRβ (#3025S), phospho-IR/IGF1R (#3024L), Akt (#4685) and phospho-Akt (S473) (#4060) from Cell Signaling Technology and goat anti-rabbit HRP conjugate (#1706515) from Bio-Rad. Protein detection was realized using a mix of a luminol solution and a peroxide solution (1:1) (Thermo Fisher Scientific). Protein bands were detected with a ChemiDoc MP Imaging System (Bio-Rad) and quantified with ImageJ.

### RNA isolation

Tissues were homogenized in 1 ml of QIAzol Lysis Reagent (Qiagen) using 0.1 mm dia Zirconia/Silica beads (Biospec) and Minibeadbeater (Biospec). After homogenization, samples were incubated for 5 min at RT, centrifuged (12000g, 10 min, 4°C) and supernatant was transferred to a new tube. In the case of BAT and WAT, an additional centrifugation step was included and the upper fatty layer was avoided when transferred to new tubes. 200 μl of chloroform (Sigma-Aldrich) was added, the samples were vortexed for 15s, incubated for 5 min at RT and centrifuged (12000g, 15 min, 4°C). The aqueous phase was transferred to new tubes, 100% ethanol in ratio 1:1 was added and subsequently the Direct-zol RNA Miniprep Kit (Zymo Research) was used according to the manufacturer’s instructions.

### RT-qPCR

DNAse treatment and cDNA synthesis was performed using the SuperScript™ IV VILO™ Master Mix with ezDNAse (Invitrogen) according to manufacturer’s instructions. The qPCR was performed using Power SYBR^®^ Green PCR Master Mix (Applied Biosystems) according to manufacturer’s instructions on QuantStudio 3 (Applied Biosystems). The primers used are listed in **Table S2**.

## Acknowledgements

This work was supported by the National Institute of Diabetes and Digestive and Kidney Diseases of the National Institutes of Health under award number K01DK117967 (to EA) and R01DK031026 and R01DK033201 (to C.R.K.). EA was also supported by The G. Harold & Leila Y. Mathers Foundation. The in vivo experiments in mice were conducted and tissues for molecular analysis were obtained from National Mouse Metabolic Phenotyping Center (MMPC) at UMass Medical School supported by an NIH grant (5U2C-DK093000 to J.K.K.). The research of TP, LZ and JJ was supported by the Medical Research Council Grant MR/R009066/1 and by the Czech Academy of Sciences Project RVO 61388963. We would like to acknowledge Xiaochen Bai for the construction of the models of dcVILPs, Qian Huang for her help with qPCR protocol and Boston College Biology Department undergraduate students Amaya Powis, Kaan Sevgi and Maxmilian Figura for their help with cell culture work.

## Author contributions

MC assisted with ITT experiments, RNA extraction, qPCR analysis and insulin signaling experiments. FM and CRK assisted with in vivo signaling and insulin tolerance test. HLN, RHF, and JKK conducted the in vivo infusion experiments in mice and obtained tissues for molecular analysis. JKK supervised the in vivo infusion experiments. LZ, JJ and TP assisted with binding competition experiments. FAV assisted with chemical synthesis of double chain VILPs. EA, MC, JJ, CRK and JKK assisted with the analysis of the data. EA and MC wrote the manuscript, while all other authors contributed. EA designed the research and supervised the project.

---

**Figure S1:**
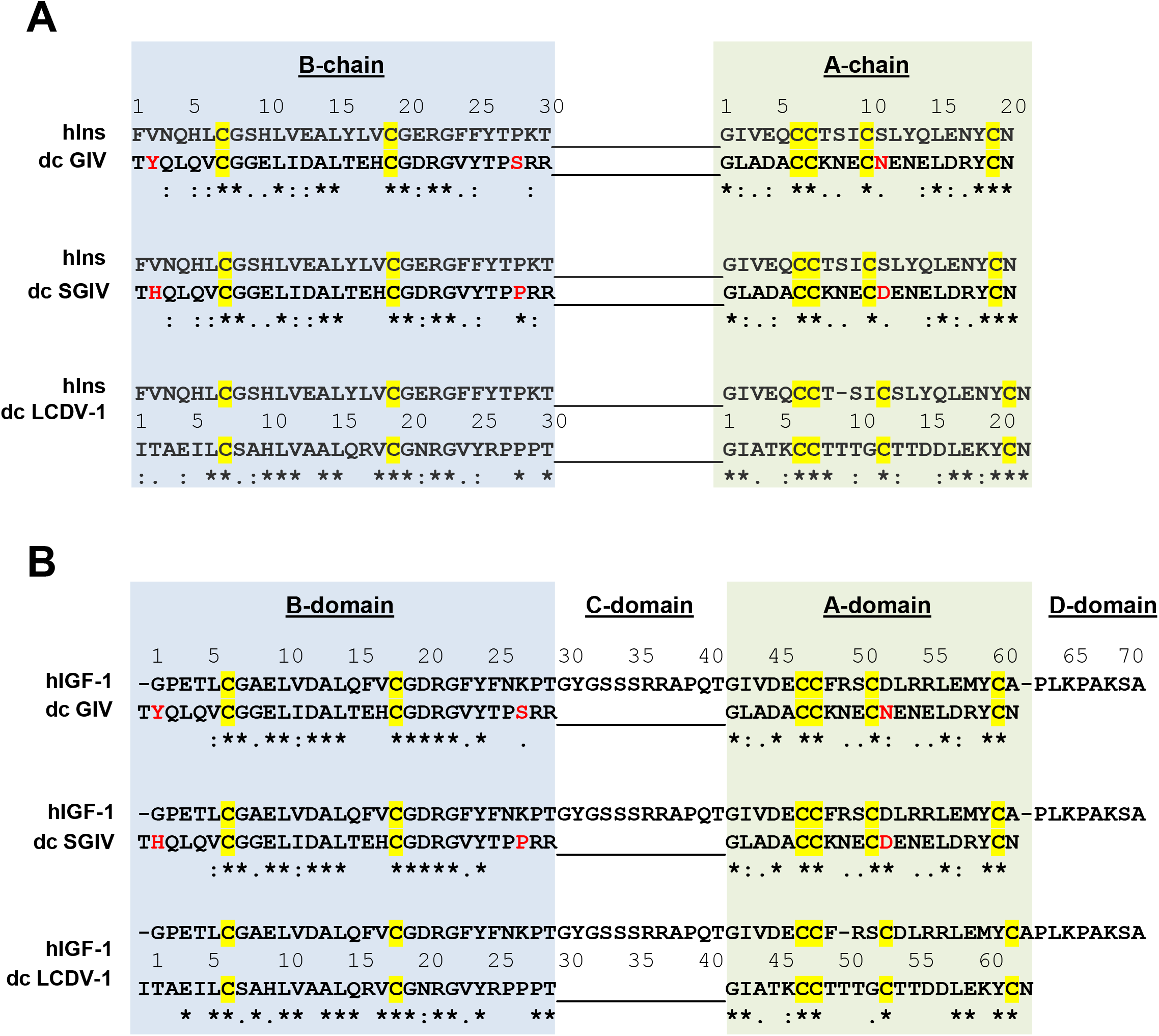
Sequence alignment of synthesized dcVILPs with human insulin and IGF-1. Comparison of dcVILPs with human insulin is shown in **A** and comparison with human IGF-1 is shown in **B**. Cysteine residues important for correct folding which are conserved in all peptides are highlighted in yellow. Three residues that differ between GIV and SGIV dcVILPs are marked in red. Conserved residues are marked by asterisk, conservatively substituted residues are marked by colon and semi-conservatively substituted residues are marked by period

**Figure S2:**
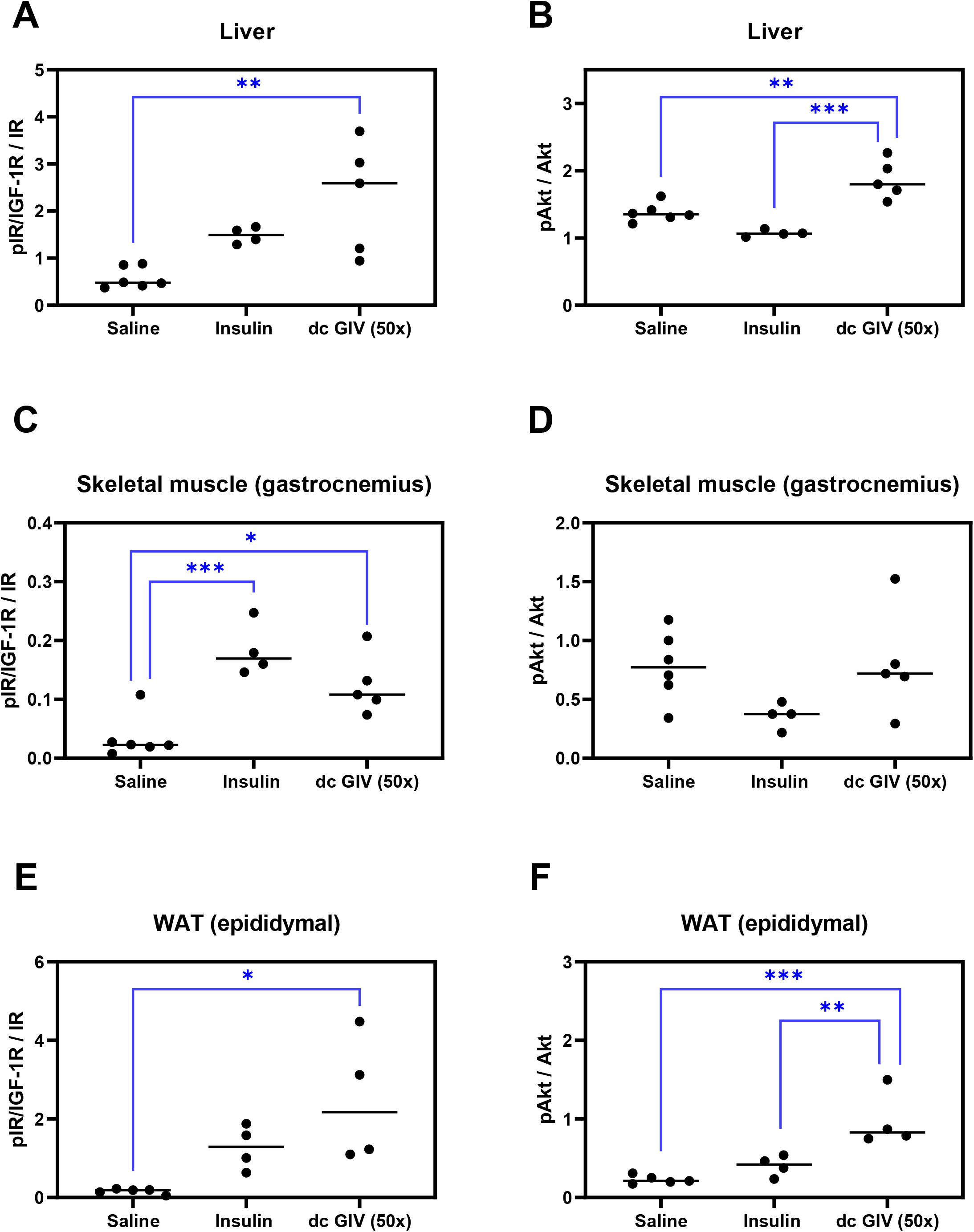
Quantification of the western blot result showing in vivo insulin signaling after in vivo experiments. Tissues were collected after 3 hours of insulin or 50x GIV VLIP infusion, and basal tissue samples were collected after 3 hours of saline infusion in awake mice. **A, C, E:** IR/IGF1R phosphorylation in liver, skeletal muscle (gastrocnemius) and WAT (epididymal), respectively. **B, D, F:** Akt phosphorylation in liver, skeletal muscle (gastrocnemius) and WAT (epididymal), respectively. n = 6 for saline, n = 4 for insulin, n = 5 for GIV dcVILP in the case of liver and skeletal muscle, n = 4 for GIV dcVILP in the case of WAT. Ordinary One-Way ANOVA followed by Tukey’s multiple comparison test was applied ((*P<0.05; **P<0.01, ***P<0.001).

**Figure S3:**
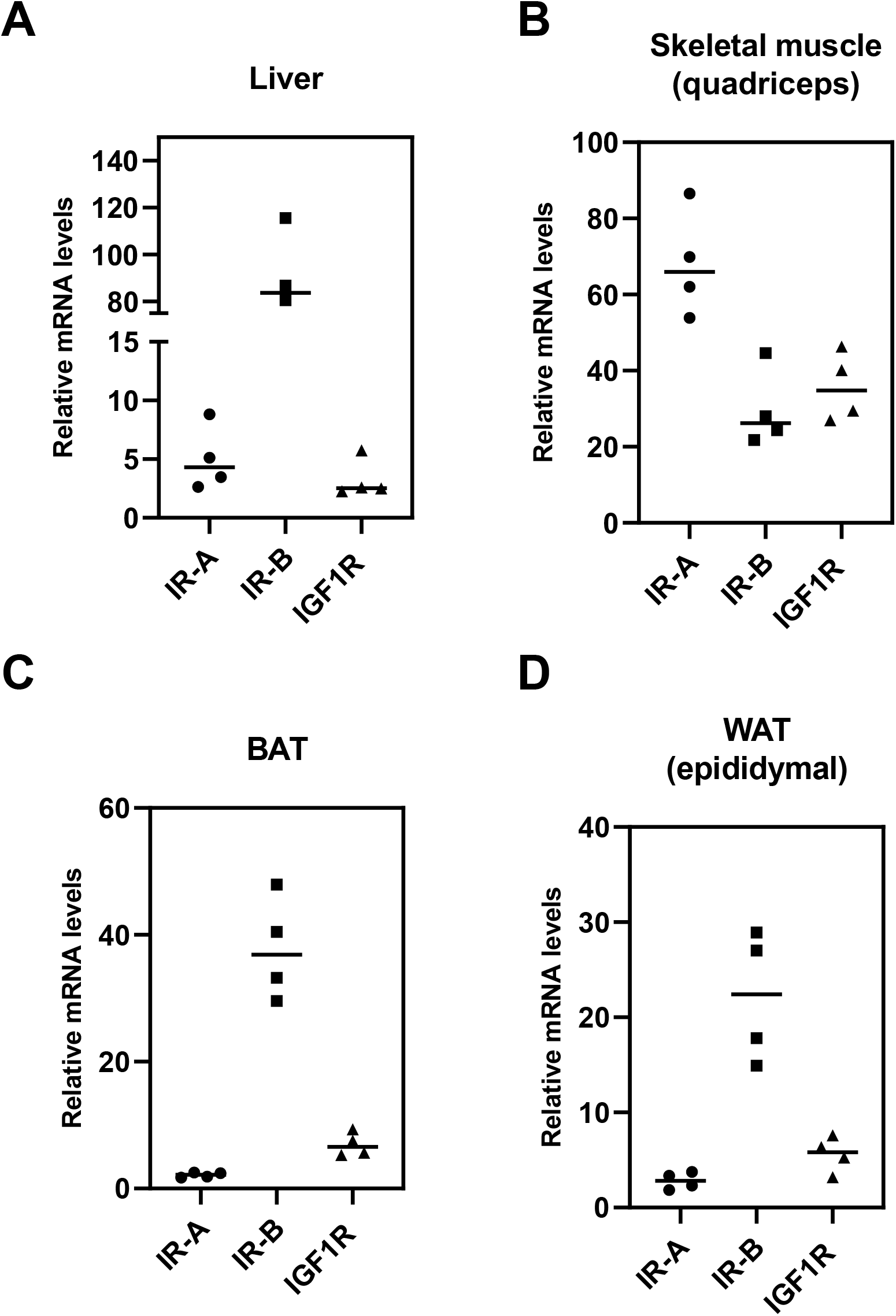
RT-qPCR analysis of IR-A, IR-B and IGF1R expression in murine liver, skeletal muscle (quadriceps), BAT and WAT. Data are expressed as % of β-actin. n = 4 per group.

**Figure S4:**
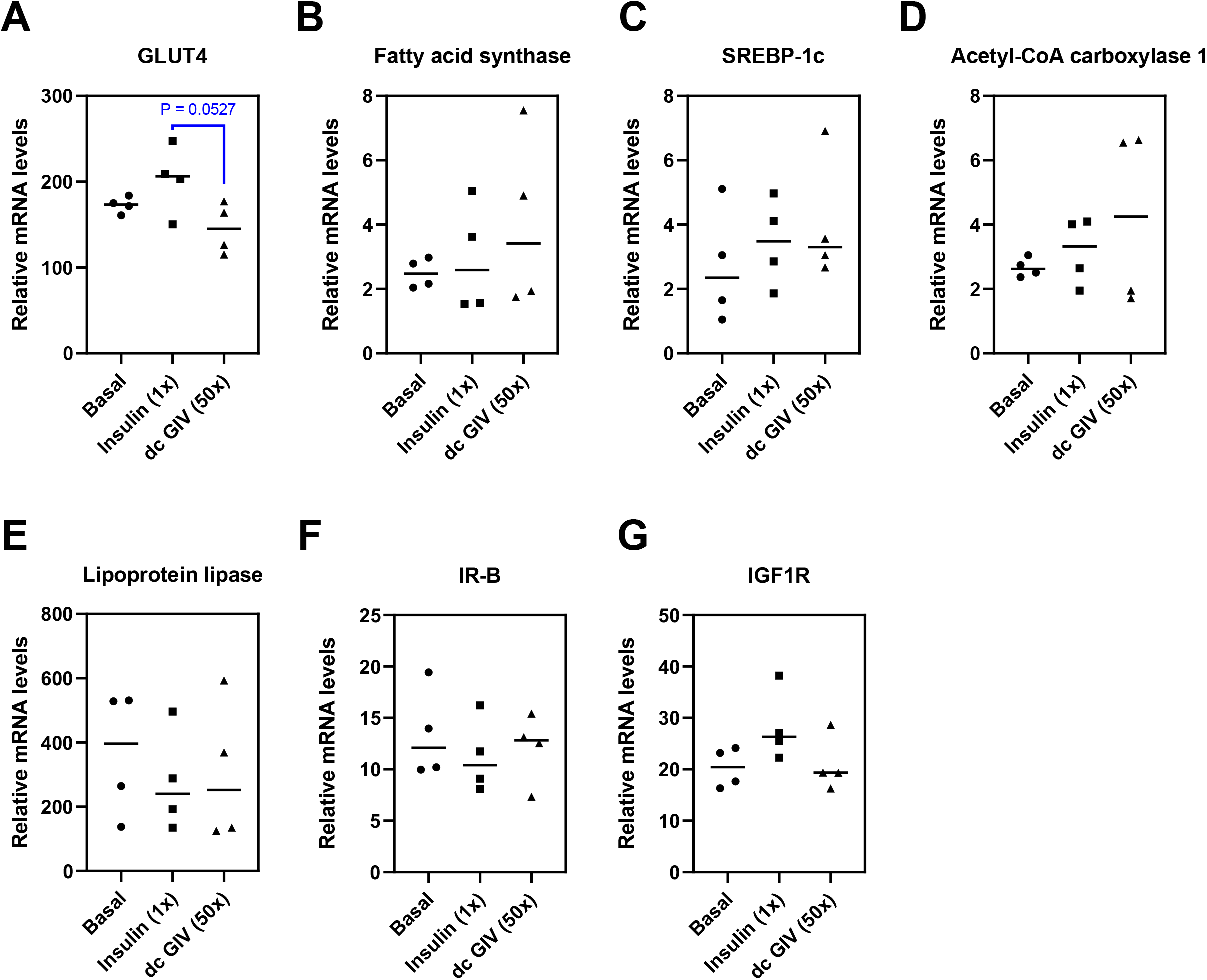
RT-qPCR analysis of expression of genes regulated by insulin action in murine skeletal muscle (quadriceps) after in vivo experiments. Tissues were collected after 3-hour insulin (0.015 nmol/kg/min; 1x) or GIV VILP (0.75 nmol/kg/min; 50x) infusion, and basal tissue samples were collected after 3 hours of saline infusion in awake mice. Data are expressed as % of β-actin. n = 4 per group. Ordinary one-way ANOVA followed by Tukey’s multiple comparison test was applied (*P<0.05; **P<0.01, ***P<0.001). P-values lower than 0.1 are indicated.

**Figure S5:**
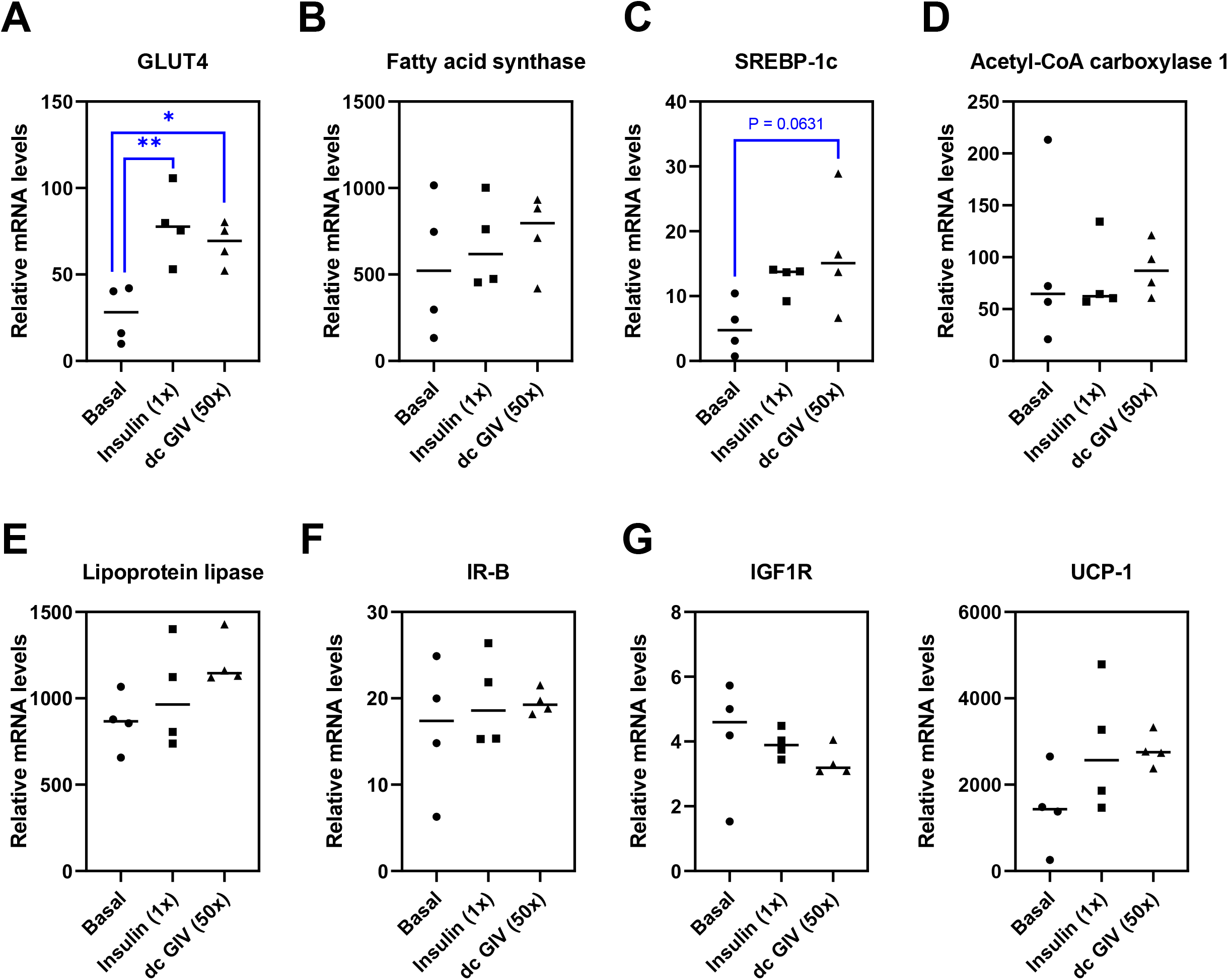
RT-qPCR analysis of expression of genes regulated by insulin action in murine BAT after in vivo experiments. Tissues were collected after 3-hour insulin (0.015 nmol/kg/min; 1x) or GIV VILP (0.75 nmol/kg/min; 50x) infusion, and basal tissue samples were collected after 3 hours of saline infusion in awake mice. Data are expressed as % of β-actin. n = 4 per group. Ordinary one-way ANOVA followed by Tukey’s multiple comparison test was applied (*P<0.05; **P<0.01, ***P<0.001). P-values lower than 0.1 are indicated.

**Figure S6:**
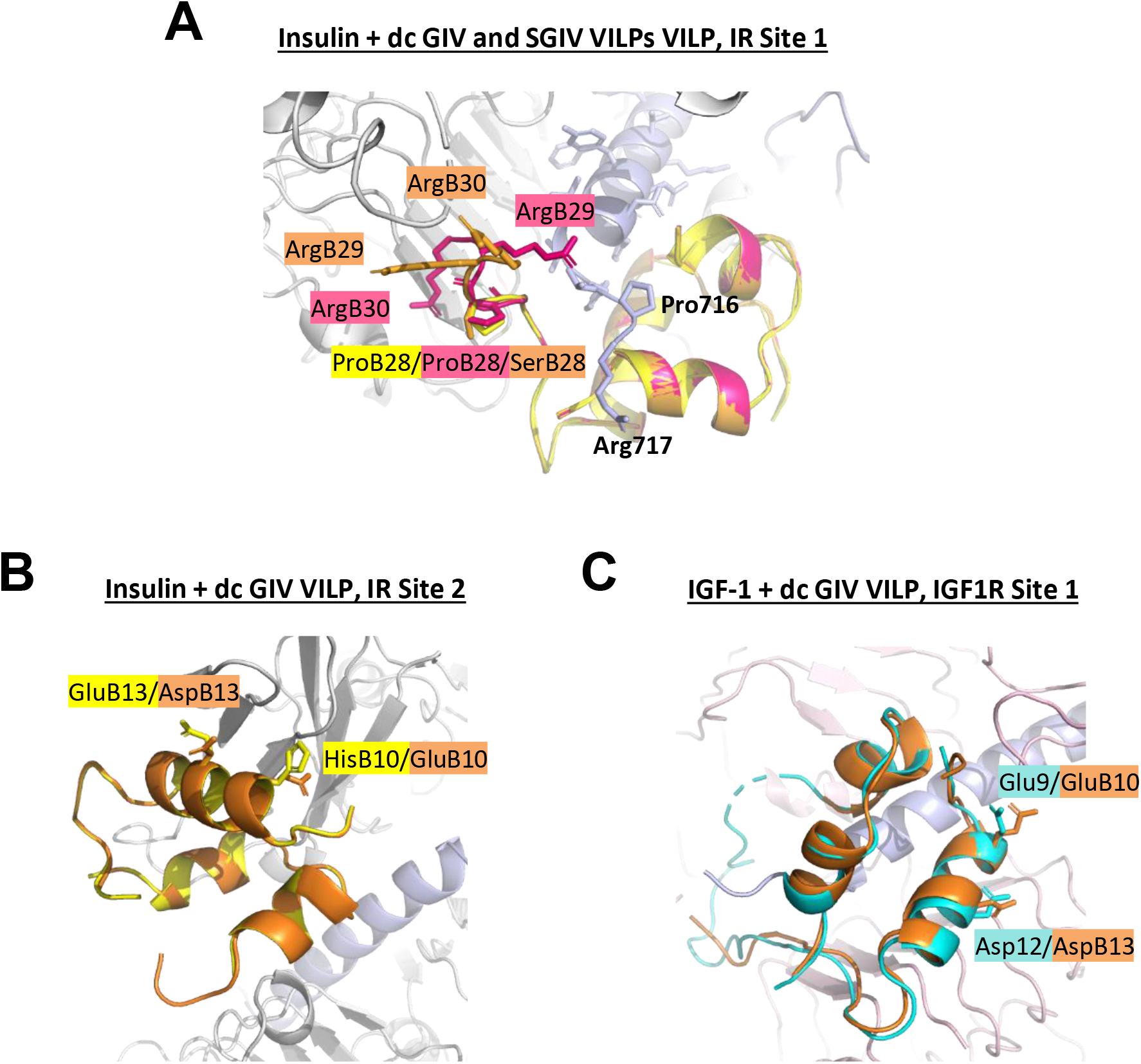
Overlay of a model of dcVILPs and insulin/IGF-1 bound to IR and IGF1R. **A:** Model of GIV and SGIV dcVILPs bound to Site 1 of IR. Insulin positions B28 (insulin and dcVILPs), B29 and B30 (dcVILPs) are shown. **B:** Model of GIV dcVILP bound to Site 2 of IR. Insulin positions GluB13 and HisB10 and their substituted counterparts in GIV dcVILP are shown. **C:** Model of GIV dcVILP bound to Site 1 of IGF1R. IGF1R positions Glu9 and Asp12 and their identical counterparts in GIV dcVILP are shown. Insulin is in yellow, IGF-1 is in magenta, GIV dcVILP is in orange and SGIV dcVILP is in pink. IR is in grey and IGF1R is in pink. The α-CT peptide is shown in light blue in all cases.

**TableS1:**
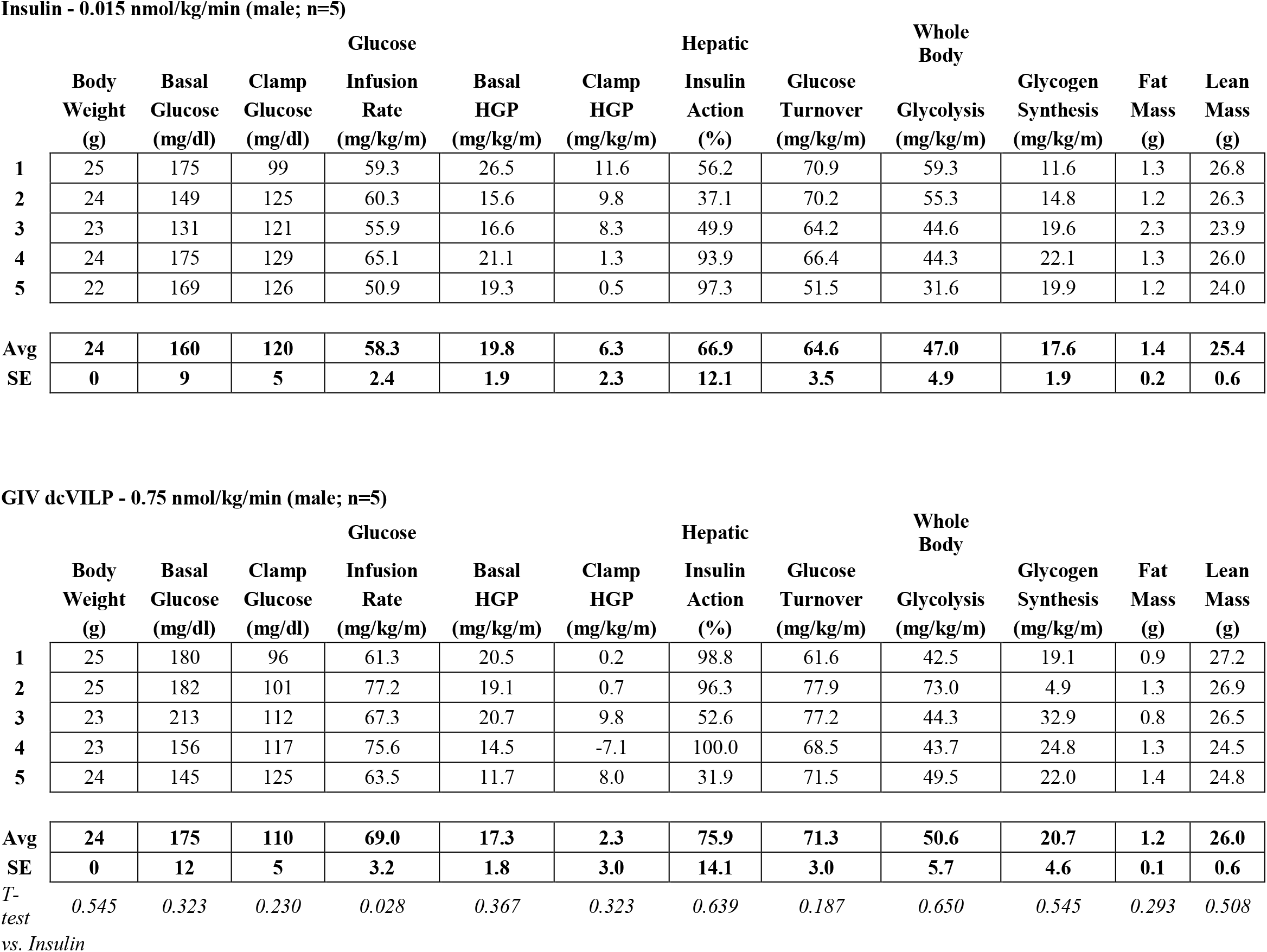
Whole body metabolism data measured in the 3-hour in vivo infusion experiments in awake mice. n = 5 for both groups.

**TableS2:**
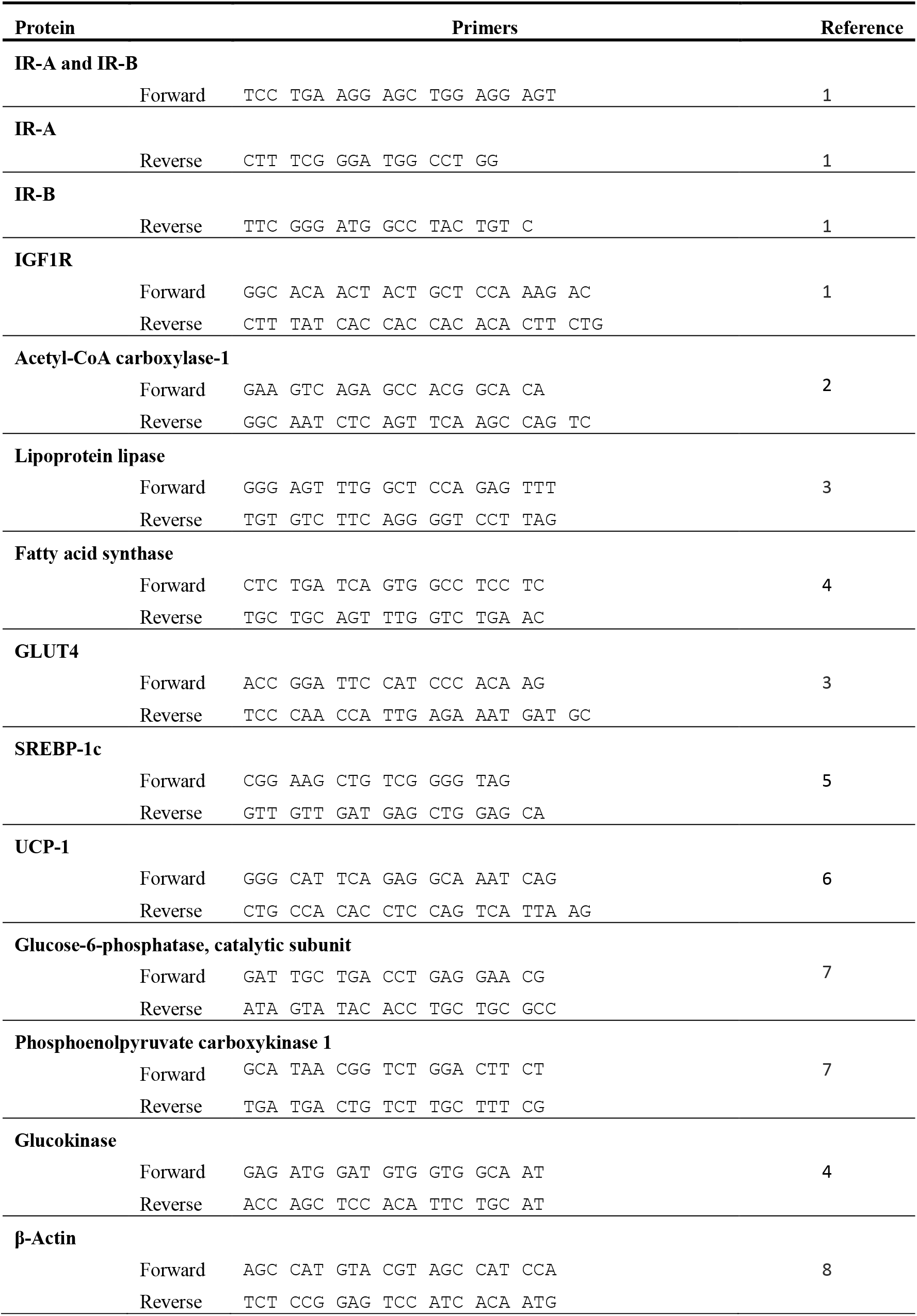
List of mouse primers used for RT-qPCR. List of primers used in this study.

